# WFS1^E864K^ in humans and mice causes Wolfram-like syndrome optic atrophy via early axonal mitochondrial dysfunction

**DOI:** 10.1101/2025.10.13.682105

**Authors:** Hernan H. Dieguez, Kevin Dubois, Elodie Reboussin, Cansu De Muijnck, Jérôme Sarniguet, Chantal Cazevieille, Stacy Alves, Julie Dégardin, Valérie Fradot, Serge Picaud, Stéphane Melik-Parsadaniantz, Maria Van Genderen, Andrea L. Vincent, Benjamin Delprat, Elodie M. Richard

**Author notes:** To whom correspondence should be addressed: Elodie M. Richard, Tel number: (33/0) 467 14 36 23, Fax number: (33/0) 467 14 92 95, Benjamin Delprat, Tel number: (33/0) 467 14 36 23, Fax number: (33/0) 467 14 92 95. **Disclosure statement:** Benjamin Delprat holds a patent on therapeutic strategy for Wolfram syndrome. No potential conflict of interest was reported by the author(s).

## Abstract

Wolfram-like syndrome leads to retinal ganglion cell degeneration and vision loss. Wolfram-like syndrome is primarily caused by variants in the *WFS1* gene, which encodes an endoplasmic reticulum resident transmembrane protein, Wolframin. To date, the disease mechanism remains unclear, and no therapies are available. Here, we generated a mouse model carrying the pathogenic *WFS1^E864K^* allele that recapitulated key features of human Wolfram-like syndrome, including bilateral optic atrophy, retinal nerve fiber thinning and lamination of the outer plexiform layer. We demonstrated, using the *Wfs1^E864K^* mouse model, that alteration of the protein leads to impairments of retinal ganglion cell function, associated with a thinning of the inner retina layer and nerve fibers. These alterations are associated with myelin disorganization, axonal death, mitochondrial alterations in the axons, and impairment of endoplasmic reticulum-mitochondria communication in the soma. Our data showed that primary deficits are localized in the optic nerve before progressing towards the retinal ganglion cell soma. RNAseq analysis identified several altered signaling pathways such as in lipid metabolism, glia activation response, metabolic stress, organelle transport and quality control. These findings highlighted the critical role of Wolframin in optic nerve mitochondrial physiology, providing us with a pertinent model to develop novel innovative therapeutic strategies.

## Introduction

Wolfram syndrome (WS) is a rare autosomal recessive disorder characterized by *diabetes insipidus*, juvenile *diabetes mellitus* (DM), optic atrophy (OA), deafness and severe neurological disabilities associated with low prognosis and premature death around 35 years of age ^1,2^. Epidemiological studies have shown a very wide frequency, ranging from 1:700,000 in the UK to 1:100,000 in North America, and 1:68,000 in the Lebanese population ^3–5^. Wolfram-like syndrome (WLS) is a dominant disorder associated with hearing impairment, OA and DM, among other symptoms ^6–9^. Both diseases are associated with variants of *WFS1* gene, which encodes a transmembrane glycoprotein of 890 amino acids known as Wolframin, localized in the endoplasmic reticulum (ER) ^10,11^. Wolframin modulates many aspects of ER and mitochondria physiology through mitochondria-associated ER membranes (MAMs) having an impact over ER stress response, calcium homeostasis, mitophagy, autophagy and mitochondrial dynamics and function, among others ^12–19^. Despite of the efforts to strike on possible therapeutic targets, as of today, no therapies exist to reverse or prevent the progression of the disease.

A precise and early diagnosis of WS and WLS pathologies are important for clinicians to ease the symptoms and enhance their prognosis. Although WLS has multiple eddects on global health, OA is the most frequent ophthalmological feature that developes during the progression of the disease (Kabanovski et al., 2022; Rigoli et al., 2022; Ustaoglu et al., 2020), with 87% of the WLS patients developing OA ^9^. In addition, some authors suggested that early detection of OA is key for a precise diagnosis of the pathology and correlates with disease severity ^22–24^. WLS causes severe visual impairment among which dyschromatopsia, visual field defects and visual acuity reduction were reported. Furthermore, retinal thickness reduction, mainly due to thinning of the inner plexiform (IPL) as well as retinal ganglion cell (RGCs) layers and retinal nerve fiber layer (RNFL) optic disk pallor and thinning of the optic nerve tracts have also been associated with WS progression ^20,25–28^. In addition to the alterations aforementioned, the lamination of the outer plexiform layer (OPL) is a key feature present only in WLS affected individuals, compared to WS affected individuals ^24,27,29^. In concordance with these ophthalmological alterations, in human retina, *WFS1* is highly expressed in the RGCs and the optic nerve (ON), similarly to what has been reported in other species such as non-human primates and mice ^30–32^.

Over the last decade, several experimental models, including human induced pluripotent cells (iPSC), fly, zebrafish, rat and mouse models, have been developed to unravel the mechanism(s) behind pathologies and explore new therapeutic strategies^33^. However, most models do not accurately replicate the pathophysiological processes of WS and fail to reproduce the correct sequence of visual alterations^14,34–40^.

Recently, we developed a murine experimental model of WLS reproducing the human variant c.2590G > A, p.(E864K). We previously showed that the *Wfs1^E864K^* homozygous mutant mice are profoundly deaf by post-natal day (P)30 with a reduction of the endocochlear potential, an atrophy of the stria vascularis and a degeneration of the neurosensory epithelium, most likely stemming from a deregulation of potassium ion transfer ^41^. In isolated patient’s fibroblasts harboring *WFS1^E864K^* allele and in cultured neurons from *Wfs1^E864K^* homozygous mutant mice, we showed a reduction of the number of MAMs, with an alteration of the mitochondria dynamics, morphology and function, a deregulation of the calcium flux, the autophagy and mitophagy processes ^12^, strengthening the hypothesis of a tissue specific role for WFS1.

In human, *WFS1^E864K^* disease-causing variant has been associated in several unrelated individuals with low-frequency sensorineural hearing loss, OA and glucose intolerance ^6,8,28,42^ but also non-syndromic low-frequency hearing loss ^43,44^. As of today, the ophthalmologic consequences of the *WFS1^E864K^* variant are not systematically assessed and documented, leaving some uncertainty in the clinical features associated with this variant. Moreover, the pathophysiological mechanism leading to the visual loss remains to be identified.

In this study, we report that *WFS1^E864K^* variant is associated with bilateral optic atrophy, retinal nerve fiber thinning and lamination of the outer plexiform layer, confirming the previously reported features of WLS optic neuropathy. Using the *Wfs1^E864K^* mouse model, we measured a decrease of the RGCs, bipolar and amacrine cells electrophysiological function as early as P30, with a progressive thinning of the inner retina layers and the outer plexiform layer, and a loss of RGCs starting at P90. The alteration of the optic nerve precedes those of the retina, with a progressive demyelination of the axons, death of nerve fibers, with no overt loss of glial cells. The exploration of mitochondria function and ultrastructure showed that mitochondria bioenergetics and structure are altered in the ON, with a decrease of ER contacts with mitochondria within the soma of the RGCs, confirming the key role of WFS1 in this organelle. Transcriptomic analyses of the ON shed light on deregulated genes suggesting, taken together with the histological and physiological analyses, a default in mitochondria transport and quality control in the ON.

## Results

### Clinical presentation of 2 unrelated affected individuals unveils the hallmarks of WLS visual alteration in WFS1^E864K^ carriers

Case NZ01 was a 10-year-old female referred to ophthalmology from ORL assessment given her history of sensorineural hearing loss. Born in Pakistan to non-consanguineous parents, the family moved to New Zealand when she was 7.5 years, and she failed school audiology screening, and subsequently determined to have bilateral severe sensorineural hearing loss. She had a history of poor speech development from age 2, with a mild microcytic hypochromic anemia, associated with Beta thalassemia trait. A magnetic resonance imagery (MRI) test showed small optic nerves, and gene testing for sensorineural hearing loss was negative. Initially she had hearing aids but was referred for cochlear implants at age 10. The initial eye assessment demonstrated a myopic astigmatism, with best corrected visual acuity (BCVA) of 0.5 for the right eye, 0.63 for the left eye, Ishihara color vision 11/13 for the right eye, 10/13 for the left eye (Supplemental Table 1), with temporal pallor of the optic nerves (Figure 1A), lamination of the outer plexiform layer on optical coherence tomography (OCT) (Figure 1B), and thinning of the retinal nerve fiber layer (Figure 1C). Gene testing identified the *WFS1^E864K^* variant, and parental segregation determined this was a *de novo* change. At last review age 11 her vision was unchanged, with a normal HBA1c, and has been lost to follow-up as the family returned to Pakistan.

**Figure 1.**
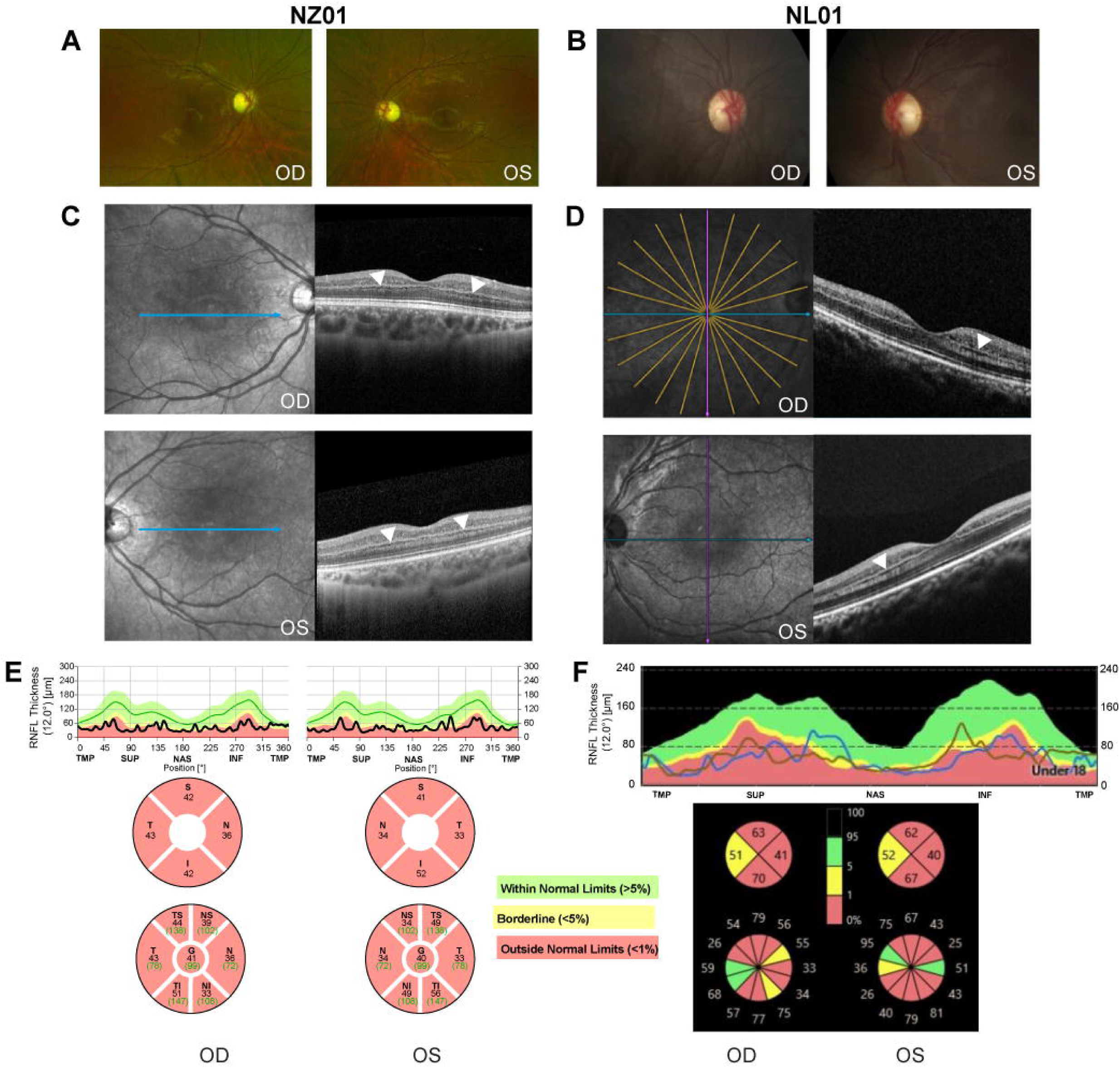
Clinical features of affected individuals NZ01 and NL01 showing a lamination of their outer plexiform layers. (A-B) Ocular fundus photographs of the right (OD) and left (OS) eyes of NZ01 (A) and NL01 (B) affected individuals, demonstrating pallor of the optic nerves, particularly temporally, but a normal retinal appearance. (C-D) Macular OCT of the right eye (OD) and left eye (OS) of both NZ01 (C) and NL01 (D). OCT reveals a lamination of the outer plexiform layer (white arrowheads) in both eyes. (E-F) Retinal nerve fiber layer (RNFL) deviation maps from the optic nerve head optical coherence tomography (OCT) confirm a significant thinning of the RNFL. The decrease of the RNFL thickness is present in all quadrants for NZ01 (E) and NL01 (F), except for the temporal quadrant in NL01. The green area corresponds to the 5^th^–95^th^ percentiles of the RNFL thickness (within normal limits) for the device reference database, the yellow area to the 1^st^–5^th^ percentiles (borderline), and the red area to below the 1^st^ percentile (outside normal limits). OD, *oculus dexter,* right eye; OS, *oculus sinister*, left eye.

Case NL01 was a 17-year-old girl who was referred for ophthalmological evaluation of optic atrophy following genetic testing for congenital hearing loss, which identified a heterozygous *WFS1^E864K^* variant. Born in South America and adopted by Dutch parents at age six, she presented with delayed language development and sensorineural hearing loss. Early assessments also revealed *pubertas precox* and low-average intellectual quotient, though language delay may have influenced cognitive testing outcomes.

She was treated with puberty-suppressing medication and initially fitted with bilateral hearing aids, later receiving bilateral cochlear implants at age 14. That same year, ophthalmologic screening revealed no visual complaints, but examination showed bilaterally pale optic discs (Figure 1B). OCT demonstrated splitting of the outer plexiform layer (Figure 1D) and significant retinal nerve fiber layer thinning (Figure 1F) — findings consistent with autosomal dominant *WFS1*-related optic neuropathy. Visual fields, color vision, and visual evoked potentials were normal (Supplemental Table 1). At her most recent visit at age 17, her visual acuity had declined slightly to 0.8 Snellen bilaterally (Supplemental Table 1), though she remained asymptomatic.

### Wfs1^E864K^ allele induces progressive retinal dysfunction in mice

To reveal the impact of the WFS1^E864K^ protein on vision, the previously described *Wfs1^E864K^* mouse model was used and all three genotypes, *Wfs1^WT^*, *Wfs1^E864K/WT^* and *Wfs1^E864K^* were studied. Retina function and histology, from both males and females, were analyzed at post-natal day (P) 30, P60 and P90 (Figure 2A). The electroretinograms (ERG) is well established tool in experimental and clinical ophthalmology to evaluate retinal function. Under scotopic conditions, positive scotopic threshold response (pSTR), oscillatory potentials (OPs), b-wave and a-wave have been studied to evaluate retinal ganglion cells (RGCs), amacrine, bipolar (OFF) cells and rods, while photopic conditions was done to explore RGCs, bipolar (ON) cells and cone function ^45–52^ The pSTR (Figure 2C), OPs (Figure 2D) and b-wave (Figure 2E) amplitudes of *Wfs1^E864K^* mice were significantly decreased compared to those of their wild-type littermates at all measured time points. In the heterozygous *Wfs1^E864K/ WT^* mice, a statistically significant decrease of the pSTR amplitude was observed as early as P30 while a decrease of the OPs and b-wave amplitudes was first measured at P60. No differences in the a-wave amplitude were recorded between groups at any time points (Figure 2F). Moreover, under a lower intensity stimulation (0.14 cd.s/m^2^), the variations of the a-wave and b-wave amplitudes (Supplemental Figure 1) were comparable to those observed under higher intensity stimulation (3.2 cd.s/m^2^) (Figure 2). Additionally, the analysis of the waveforms recorded under photopic conditions (Supplemental Figure 2A), showed a decrease of both photopic b-wave (Supplemental Figure 2B) and phNR (Supplemental Figure 2C) amplitudes in *Wfs1^E864K^* mice, at all time points, compared to their wild-type and heterozygous littermates.

**Figure 2.**
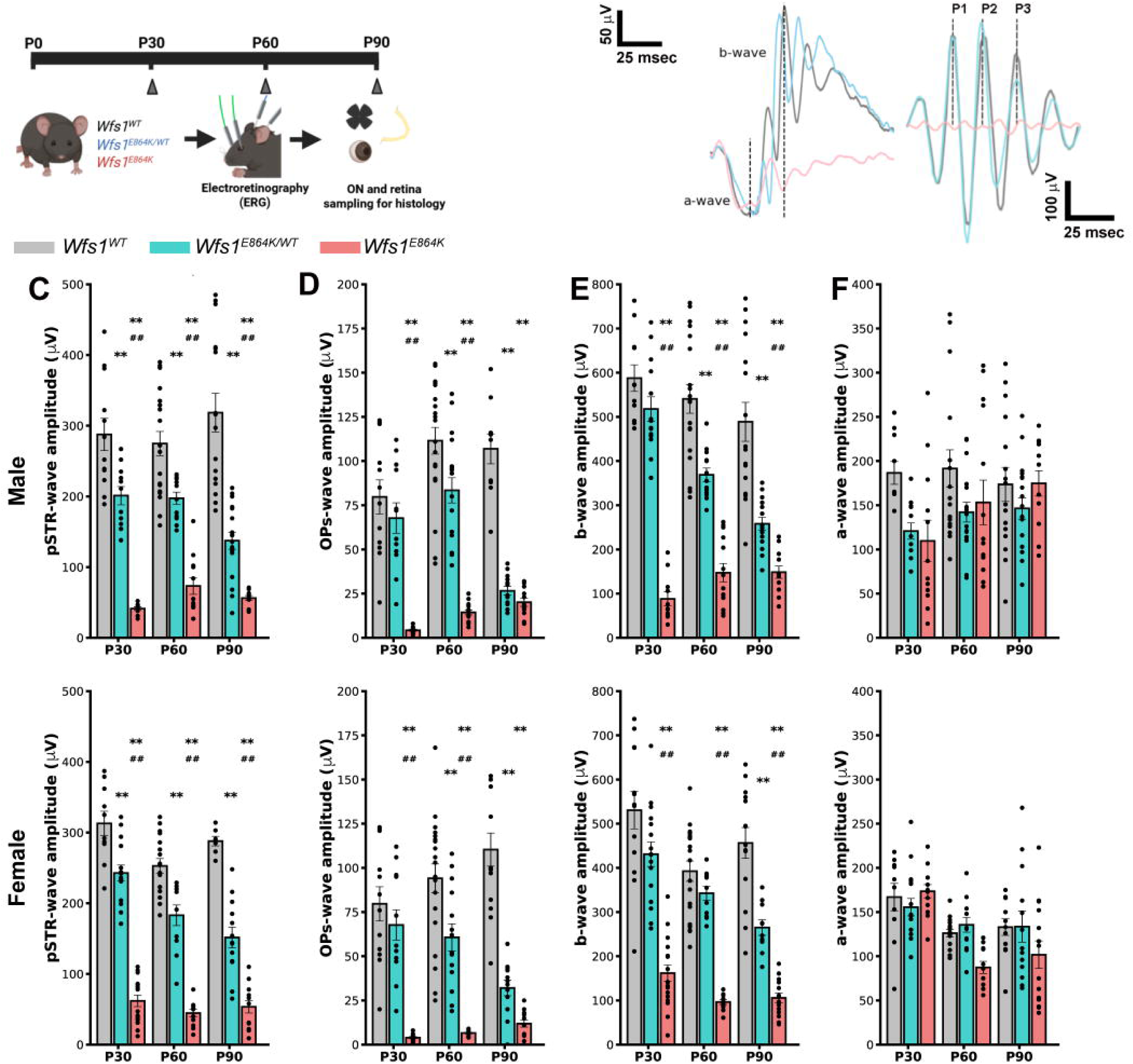
*Wfs1^E864K^* allele is associated with scotopic retinal dysfunction. (A) Representative schematic of the experimental procedure. Retina function of *Wfs1^WT^*, *Wfs1^E864K/WT^* and *Wfs1^E864K^* mice were assessed by ERG at P30, P60 and P90. Retinas and optic nerves (ONs) were collected for histological analysis at each time point. (B) Representative scotopic ERG traces showing pSTR amplitude at 0.14 mcd.s/m^2^ (left), a-wave and b-wave amplitudes at 3.2 mcd.s/m^2^ (middle) and OPs waves at at 1.2 mcd.s/m^2^ (right) from *Wfs1^WT^* (grey), *Wfs1^E864K/WT^* (blue) and *Wfs1^E864K^* (red) male mice at P30. (C-F) Quantification of pSTR (C), OPs (D), b-wave (E), and a-wave amplitude (F) for *Wfs1^WT^* (grey), *Wfs1^E864K/WT^* (blue) and *Wfs1^E864K^* (red) mice at P30, P60 P90, for both males and females. Data are presented as mean ± SEM from both eyes of 5 to 10 mice. One-way Anova test was performed for each time point, followed by a post-hoc Tukey’s test. ** p < 0.01 vs. *Wfs1^WT^* mice, ## p < 0.01 vs. *Wfs1^E864K/WT^* mice. ERG: electroretinograms; ON: optic nerve; pSTR: positive scotopic threshold response; OPs: oscillary potentials.

### A thinning of the retinal inner layers and the outer plexiform layer is observed in homozygous Wfs1^E864K^ mice, with signs of apoptosis

As our electrophysiological recordings suggest a dysfunction in the inner layers of the retina, as early as P30, we performed hematoxylin and eosin staining on transversal retinal sections of all genotype and age, to evaluate the damage at the organ level (Figure 3A). At P90, a decrease of the thickness of the retina of *Wfs1^E864K^* mice, compared to their heterozygous and wild-type littermates, was measured (Figure 3B). To pinpoint which cell type(s) might be affected, we measured the thickness of each retinal layer. Although, no changes were found in the outer segment (OS) nor outer nuclear layers (ONL) at any of the measured time points (Figure 3C and 3D), a progressive thinning of the outer plexiform layer (OPL), starting at P30, was found in *Wfs1^E864K^* mice (Figure 3E). In addition, inner nuclear layer (INL) and inner plexiform layer (IPL) thickness were significantly decreased at P90 in the *Wfs1^E864K^* mice compared to the wild-type controls (Figure 3F and 3G). In the heterozygous *Wfs1^E864K/WT^* mice, thinner OPL and IPL, compared to control littermates were measured at P90 (Figure 3E and 3G).

**Figure 3.**
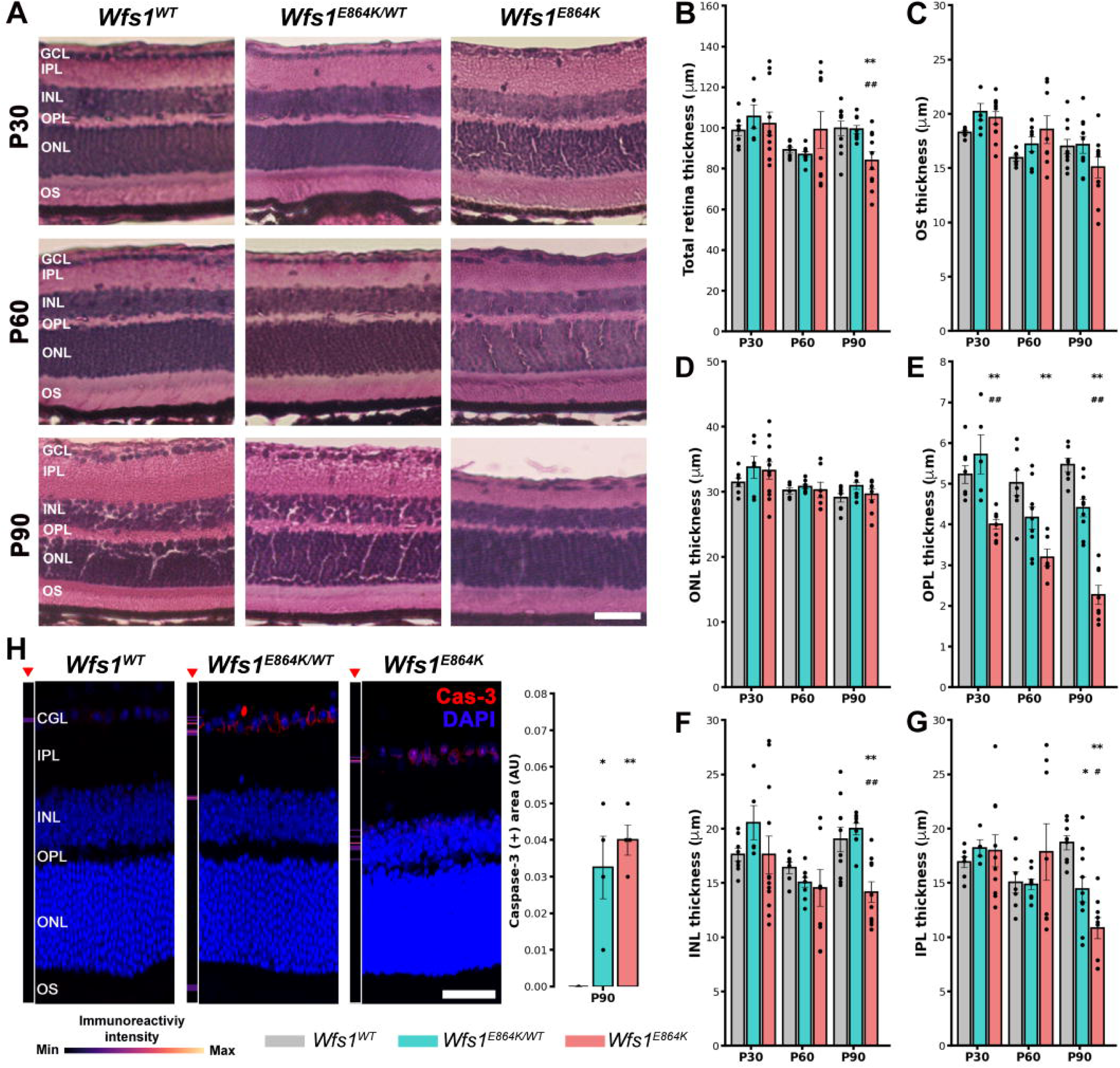
*Wfs1^E864K^* mutant mice develop retinal morphometry alterations, due to apoptosis in a dose dependent fashion. (A) Representative images of retina paraffin transversal sections following a hematoxylin and eosin staining from *Wfs1^WT^*, *Wfs1^E864K/WT^* and *Wfs1^E864K^* mice at P30, P60 and P90. (B-G) Quantification of total retina (B), outer segment (OS) (C), outer nuclear layer (ONL) (D), outer plexiform layer (OPL) (E), inner nuclear layer (INL) (F) and inner plexiform layer (IPL) (G) thickness in *Wfs1^WT^*, *Wfs1^E864K/WT^* and *Wfs1^E864K^* mice at P30, P60 and P90. (H) Representative confocal images of immunostaining for cleaved caspase-3 (red) on retina transversal sections, counterstained with DAPI (blue), with a heatmap immunoreactive transversal analysis (red head arrow) and quantification of caspase-3 positive area thickness for *Wfs1^WT^* (grey), *Wfs1^E864K/WT^* (blue) and *Wfs1^E864K^* (red) mice at P90. Data are presented as mean ± SEM from 5 to 10 retina sections. Scale bar: 30 mm for all panels in (A) and 20mm for all panels in (H). One-way Anova test was performed for each time point, followed by a post-hoc Tukey’s test. * p < 0.05, ** p < 0.01 vs. *Wfs1^WT^* mice, # p < 0.05, ## p < 0.01 vs. *Wfs1^E864K/WT^* mice.

Additionally, to confirm if the retina morphometry changes could be associated to neurodegeneration and apoptosis, immunolabeling with cleaved caspase-3 antibodies was performed at P90 (Figure 3H). The quantification of the immunoreactivity positive area has unveiled the presence of apoptosis in both the homozygous and heterozygous *Wfs1^E864K^* mutant mice, localized predominantly in the inner retina, as highlighted by the heatmap analysis across the retina transversal section (red arrow, Figure 3H).

### Wfs1^E864K^ aberrant protein affects bipolar cells and RGCs, with an increase of the retina reactive gliosis

The retina being composed of various cell type layers, to better explore which cell types are affected by *Wfs1^E864K^* variant, immunolabeling of transversal retina sections were performed with the following antibodies: protein kinase C one alpha (PKC-1a) and synaptophysin to highlight the bipolar cells and their synapsis, Glial fibrillar acid protein (GFAP) for retina reactive gliosis, Rhodopsin for rods and cone Arrestin for cones. Although, the *Wfs1^E864K^* allele did not induce any significant alteration at P30, it induced a significant decrease in PKC-1a, Synaptophysin and cone Arrestin immunoreactivity positive area only in *Wfs1^E864K^* mice at P60 and P90 (Figure 4A, 4B and Supplemental Figure 3A and 3B) along with a significant increase in GFAP positive area at P60, predominantly in the inner retina (Figure 4C). Notably, Rhodopsin positive area did not change across time in any genotypes (Supplemental Figure 3C and 3D).

**Figure 4.**
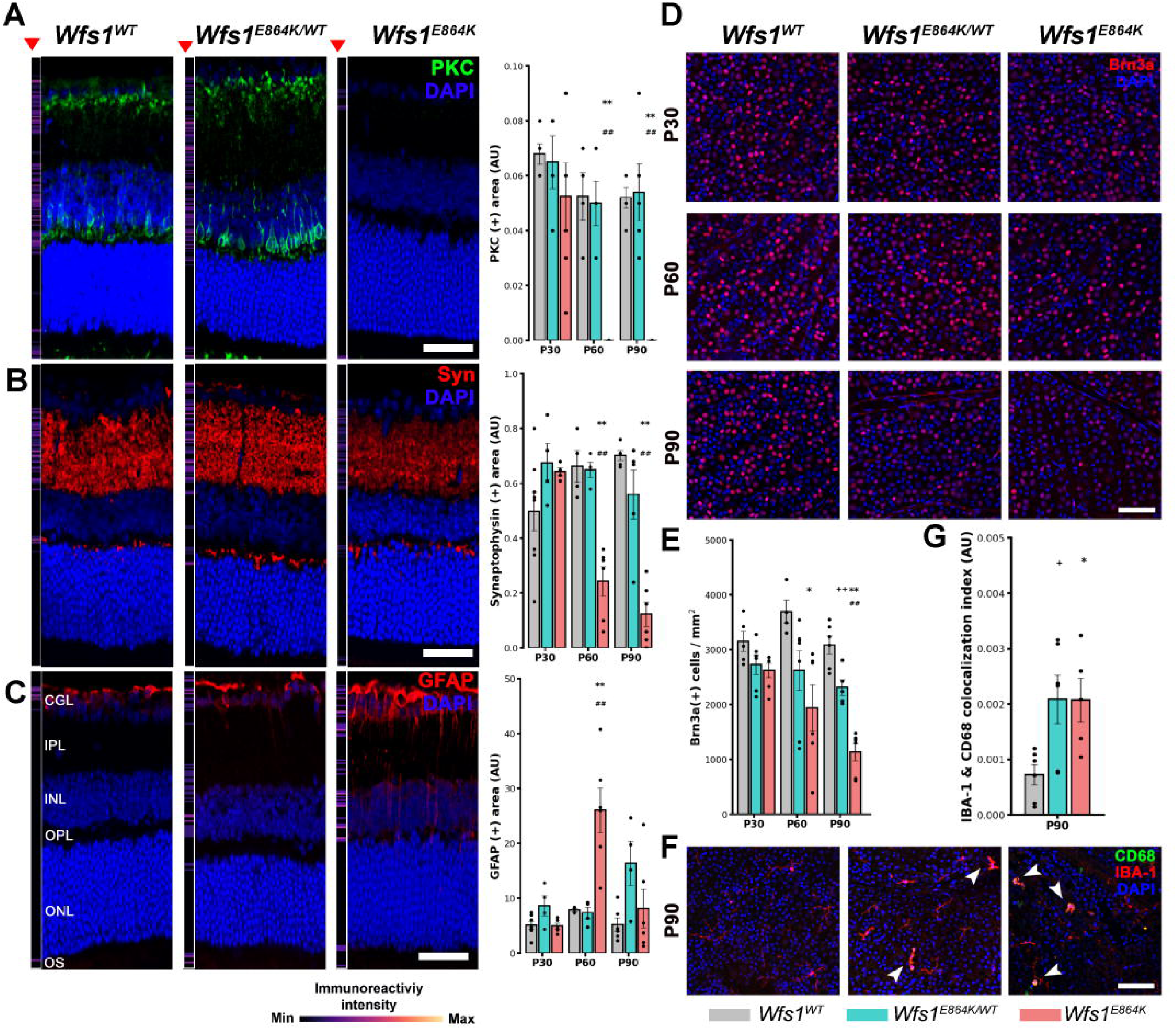
Bipolar cells and RGCs are affected in *Wfs1^E864K^* mutant mice. (A - C) Representative confocal images of immunostaining with PKC-1a (A, green), synaptophysin (Syn) (B, red) and GFAP (C, red) antibodies on transversal sections of *Wfs1^WT^*, *Wfs1^E864K/WT^* and *Wfs1^E864K^* mice at P60 and associated heatmap immunoreactive transversal analysis (red head arrow), alongside quantification of immunoreactivity positive area for each antibody in *Wfs1^WT^* (grey), *Wfs1^E864K/WT^* (blue) and *Wfs1^E864K^* (red) mice at P30, P60 and P90. (D-E) Representative confocal images of immunostaining with Brn3a (red) on retina flat mounts, counterstained with DAPI (blue) from *Wfs1^WT^* (grey), *Wfs1^E864K/WT^* (blue) and *Wfs1^E864K^* (red) mice at P30, P60 and P90 (D) and the quantification of the number of Brn3a positive cells per mm^2^ per condition (E). (F) Representative confocal images of immunostaining with IBA-1(green) and CD68 (red) antibodies on retina flat mounts, counterstained with DAPI (blue) from *Wfs1^WT^* (grey), *Wfs1^E864K/WT^* (blue) and *Wfs1^E864K^* (red) mice at P90 (F) and the quantification of the number of IBA-1 and CD68 positive cells per image (F). Data are presented as mean ± SEM from 5 to 6 retina sections and 5 to 7 retina flat mounts. Scale bar: 20 mm for all panels in (A-C), 50mm for all panels in (D) and (F). One-way Anova test was performed for each time point, followed by a post-hoc Tukey’s test. * p < 0.05, ** p < 0.01 vs. *Wfs1^WT^* mice, # p < 0.05, ## p < 0.01 vs. *Wfs1^E864K/WT^* mice.

Considering that RGCs loss and RNFL defects are key features of WLS affected individuals’ visual deficits, as shown in Figure 1, we immunolabeled RGCs with Brn3a antibody in retinal flat mounts (Figure 4D). The number of Brn3a positive cells/mm^2^ was significantly decreased as early as P60 in *Wfs1^E864K^* mice, and at P90 in *Wfs1^E864K/ WT^* mice (Figure 4E). Moreover, IBA-1 and CD68 immunostainings were performed to study microglia pro-inflammatory activation and macrophages, respectively, at our latest time point (P90) (Figure 4F). A significant increase of IBA-1 and CD68 colocalization index was found in both *Wfs1^E864K^* variant mice in the RGC layer (Figure 4G).

### Optic atrophy is due to both myelination defects and axonal loss in Wfs1^E864K^ heterozygous and homozygous mutant mice

In human, optic nerve atrophy is associated with *WFS1^E864K^* allele as shown in Figure 1. The gold-standard assay for describing ON damage in mice is to quantify the number of axons and their level of myelination. Therefore, we collected ON from all three genotypes were collected for histological processing at P30, P60 and P90.

A thinning of the optic nerve diameter was measured in *Wfs1^E864K^* mice, in semi-thin ON sections stained with toluidine, at P90 (Figure 5A and 5B). To determine the cause of this thinning, we counted the number of axons per ON section, on transmitted electron micrographs, and put in light a decrease in *Wfs1^E864K^* mice (Figure 5A and C). The axons were then classified, according to their size (small: 0.3 - 0.8 mm, medium: 0.8 – 1.4 mm, and large: 1.4 – 2.2 mm). Thus, the smaller number of axons in *Wfs1^E864K^* homozygous mice could be explained by the decrease of the number of large and medium-sized axons. Conversely, the number of large-sized axons was decreased in *Wfs1^E864K/WT^* mice without affecting significantly the total number of axons (Figure 5D), in line with the unchanged diameter of the ON in these mice.

**Figure 5.**
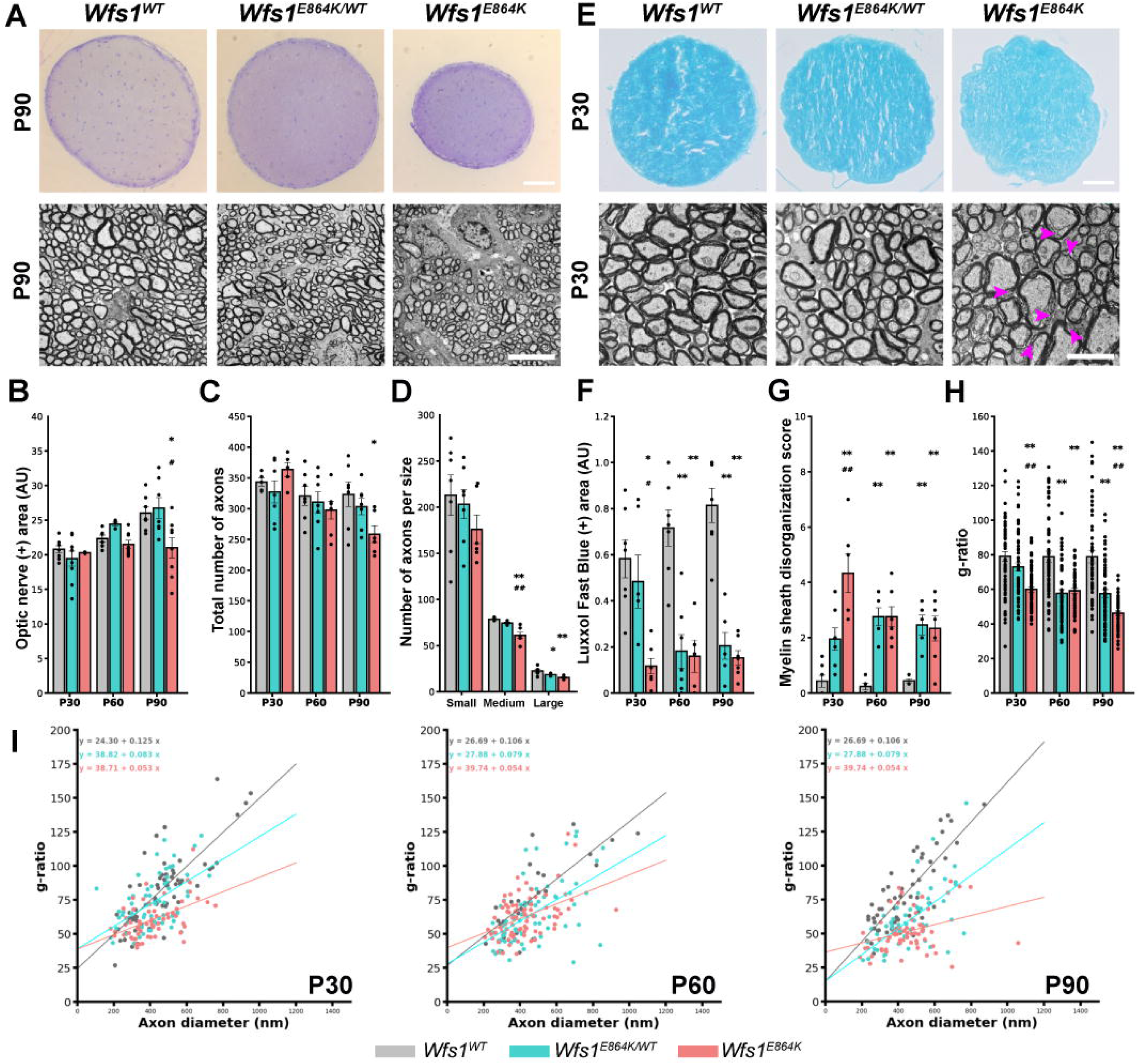
Wfs1^E864K^ protein is leading to early optic nerve atrophy in both heterozygous and homozygous mutant mice. (A) Representative images of semi-thin optic nerve sections stained with toluidine (upper panel) and transmission electron micrographs of the transversal ON (lower panel) of *Wfs1^WT^*, *Wfs1^E864K/WT^* and *Wfs1^E864K^* mice at P90. (B - D) Quantification of ON relative area (B), total number of axons (C) and number of axons per size of *Wfs1^WT^* (grey), *Wfs1^E864K/WT^* (blue) and *Wfs1^E864K^* (red) mice at P30, P60 and P90. (E) Representative images of ON transversal sections stained with Luxol Fast Blue (LFB) (upper panel) and transmission electron micrographs of the transversal ON (lower panel) from *Wfs1^WT^*, *Wfs1^E864K/WT^* and *Wfs1^E864K^* mice at P30. Magenta arrows highlight disorganized myelin sheath in lower panel (E). (F-H) Quantification of LFB positive area (F), myelin sheath disorganization score (G) and myelin g-ratio (H) of *Wfs1^WT^* (grey), *Wfs1^E864K/WT^* (blue) and *Wfs1^E864K^* (red) mice at P30, P60 and P90. (I) Association between g-ratio and axon diameter for *Wfs1^WT^* (grey), *Wfs1^E864K/WT^* (blue) and *Wfs1^E864K^* (red) mice at P30, P60 and P90. Data are presented as mean ± SEM from 5 to 7 ONs. Scale bars 60 mm for all panels in top (A), 2 mm for all panels in bottom (A), 60 mm for all panels in top (E) and 500 nm for all panels in bottom (E). One-way Anova test was performed for each time point, followed by a post-hoc Tukey’s test in (B-I). A Spearman correlation test was performed, followed by an ANCOVA analysis in (H). * p < 0.05, ** p < 0.01 vs. *Wfs1^WT^* mice, # p < 0.05, ## p < 0.01 vs. *Wfs1^E864K/WT^* mice.

In addition to the number of axons, their myelination status could explain not only the thinning of the ON diameter but also the electrophysiological deficits measured. Therefore, the myelin content was studied by LFB staining (Figure 5E). Notably, a progressive demyelination was observed, in a *Wfs1^E864K^* allele dose-dependent fashion, starting at P30 in the homozygous mutant mice and at a later time point, P60, in the heterozygous mice (Figure 5F). To look further into the myelination process, we used transmission electron microscopy to assess the ultrastructure of the myelin sheaths at P30 (Figure 5E), using a scoring scale detailed in the methods section to quantify the myelin sheath disorganization in all conditions. In parallel, we calculated the g-ratio (the ratio between the inner and the outer diameter of the myelin sheath), a well-established index of axonal myelination and integrity. Concomitant to an enhanced myelin sheath disorganization in *Wfs1^E864K^* mutant mice (Figure 5G), the g-ratio decreased, starting at P30 in *Wfs1^E864K^* and at P60 in *Wfs1^E864K/ WT^* mice (Figure 5H). Moreover, the relation between g-ratio and axon diameter showed significant differences between *Wfs1^E864K^* mice, with a lower R coefficient in *Wfs1^E864K^* mice at all measured time points, as early as P30 (Figure 5I), suggesting a progressive deterioration of the myelin sheath over time.

Others have reported a chronic neuroinflammatory state in astrocytes and microglia of the optic nerve in a Wfs1 knock-out mouse model, *Wfs1^exon8del^* ^36^. Therefore, we labeled the neuronal and glial cells in transversal ON sections of P30 mice of all three genotypes using the following antibodies: b-tubulin 3 (TUJ1) for RGCs axons, GFAP for astrocytes, IBA-1 for microglia and Oligo2 for oligodendrocytes (Supplemental Figure 4A). The decrease of TUJ1 positive area, reflecting the neuronal cell area, in both *Wfs1^E864K^* mutant mice, was in line with what we showed in electron transmitted micrographs, confirming a nerve fiber loss (Supplemental Figure 4B). An increase of the GFAP positive area was measured in the homozygous mice, with only a trend in the heterozygous ones (Supplemental Figure 4C), suggesting inflammatory state, with no changes in the other glial cell populations (Supplemental Figure 4D and 4E).

### Mitochondria structure and function are altered in Wfs1^E864K^ mouse optic nerve

The links between mitochondria and WFS1 have been well-established in the past decade, not only due to its localization at the MAMs but also as deregulation of WFS1 leads to abnormal mitochondrial homeostasis ^16,38,40,53,54^. More particularly, we showed that, in the context of *WFS1^E84K^* WLS, this nonsynonymous variant induces a deregulation of the mitochondrial bioenergetics with a concomitant decrease of MAMs ^12^. Therefore, considering that the ON is the primary area affected by *Wfs1^E864K^* variant in this mouse model, we investigated the mitochondrial respiration in ON explants of *Wfs1^WT^*, *Wfs1^E864K/WT^* and *Wfs1^E864K^* mice, at P30, using a Seahorse XFe analyzer. The measurement of the oxygen consumption rate (OCR) did not unveil any differences between groups, under basal conditions, in the basal respiration, the non-proton leak, the ATP production, the maximal nor the non-mitochondrial respiration (Figure 6A and 6B). Taking into account that optic nerve is a mixed cell population with both neuronal and glial cells, we incubated the explants with 2,5 mM of fluorocitrate (Flc) solution, an inhibitor of mitochondria metabolism in glial cells ^55,56^, to measure the respiration of axons only. As a control of the fluorocitrate effect, we calculated the percentage of OCR change between Flc and basal conditions, validating our protocol with a nearly 25% reduction of the OCR after Flc incubation, for each group (Supplemental Figure 5). Remarkedly, a strong disruption of the mitochondria bioenergetics was highlighted with a reduction of all parameters in *Wfs1^E864K^* ON explants, while only the basal respiration and maximal respiratory rate were reduced in *Wfs1^E864K/WT^* mice, showing a dose dependent impact of the Wfs1^E864K^ protein (Figure 6C).

**Figure 6.**
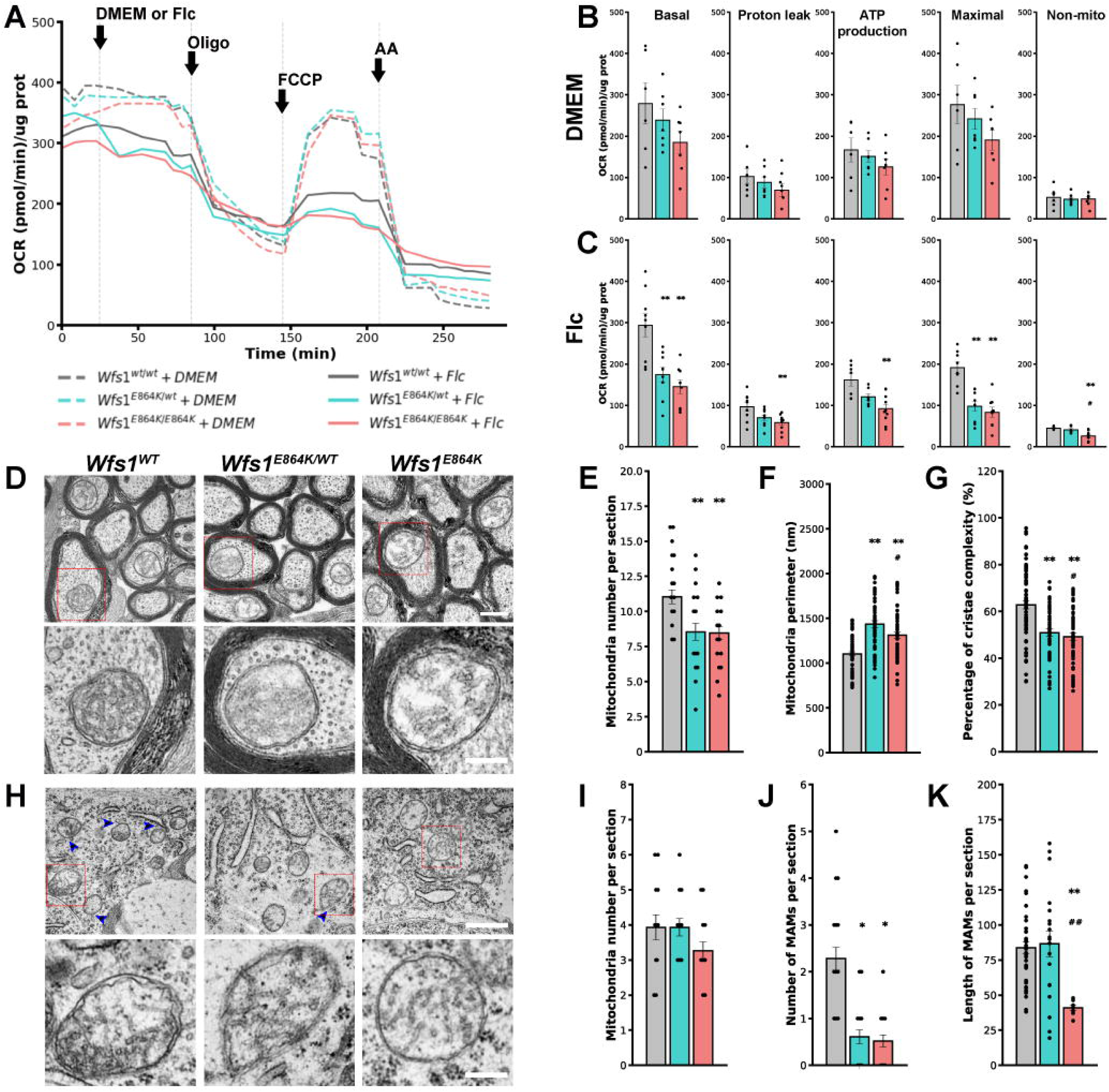
Axonal mitochondria are impaired in *Wfs1^E864K^* mice. (A) Representative traces for oxygen consumption rates (OCR) of ONs isolated from *Wfs1^WT^* (gray), *Wfs1^E864K/WT^* (blue) and *Wfs1^E864K^* (red) mice at P30, expressed as picomoles of O_2_ per minute per mg protein under basal conditions and after Oligomycin (100 mg/ml), FCCP (44 mM), Rot/AA (130 mM), under standard condition (DM; dashed line) or after injection of 2,5 mM of Fluorocitrate (Flc; plain line). (B-C) Quantification of the basal, proton leak, ATP production, maximal and non-mitochondrial respiration, calculated from OCR traces of ON from *Wfs1^WT^* (gray), *Wfs1^E864K/WT^* (blue) and *Wfs1^E864K^* (red), incubated with DM (B) or Flc (C). (D) Representative transmission electron micrographs of the mitochondria ultrastructure from the transversal ON. The lower micrographs are zoom-in of the red boxed area in the upper micrographs. (F) from *Wfs1^WT^*, *Wfs1^E864K/WT^* and *Wfs1^E864K^* mice at P30 (). (E-G) Quantification of the number (E), perimeter (F) and cristae complexity score (G) of *Wfs1^WT^* (gray), *Wfs1^E864K/WT^* (blue) and *Wfs1^E864K^* (red) mice mitochondria at P30. (H) Representative transmission electron micrographs of the mitochondria ultrastructure and of MAMs from RGCs transversal retina sections. The lower micrographs are zoom-in of the red boxed area in the upper micrographs. Blue arrows are highlighting the MAMs. (I-K) Quantification of the number of mitonchondria (I), number (J) and length (K) of MAMs from RGCs of *Wfs1^WT^* (gray), *Wfs1^E864K/WT^* (blue) and *Wfs1^E864K^* (red) mice at P30. Quantification of the mitochondria perimeter, cristae complexity score, number and length of MAMs of *Wfs1^WT/WT^*, *Wfs1^WT/E864K^* and *Wfs1^E864K/E864K^* mice at P30. Data are presented as mean ± SEM values. Data were acquired from both ONs of 6 to 8 mice for (A-C), 70 – 50 mitochondria from 5 to 7 ONs and 100 – 75 mitochondria from 5 retinas for mitochondria morphology and MAMs analysis. Scale bars: 400 nm for all images in upper panel (D), 250 nm for all images in lower panels (D) and (H), and 1000 nm for all images in upper panel (H). One-way Anova test was performed for each time point, followed by a post-hoc Tukey’s test. ** p < 0.01 vs. *Wfs1^WT^* mice, # p < 0.05, ## p < 0.01 vs. *Wfs1^E864K/WT^* mice. Oligo, oligomycin; FCCP, carbonyl cyanide-4(trifluoromethoxy)phenylhydrazone; Rot/AA, rotenone+antimycin A; DMEM, standard culture medium; Flc, fluorocitrate.

Previous studies have established a link between the bioenergetic deregulation of mitochondria, their number and structure ^12,40,54^. Therefore, in transversal ON semi-thin sections, we looked closely at the mitochondria and observed a decrease of their number in the axons of *Wfs1^E864K^* and *Wfs1^E864K/WT^* mice at P30, compared to the wild-type controls (Figure 6D and 6E). In addition, the analysis of their ultrastructure put in light a significant increase of the mitochondria perimeter (Figure 6F), a decrease of the cristae complexity in both heterozygous and homozygous *Wfs1^E864K^* mutant mice (Figure 6G), with no differences between groups for the mitochondria elongation factor (Supplemental Figure 6).

We then looked at the mitochondria at the RGC level, to assess if these defects were also seen in the soma. The retinas from all three genotypes were collected at P30 and processed for transmitted electron microscopy. Curiously, the mitochondria looked healthy in all 3 genotypes, with no variation of the elongation factor, perimeter nor cristae complexity (Supplemental Figure 6). Despite similar shape characteristics compared to those of the wild-type’s, we observed a reduction of the mitochondria contact with the ER (Figure 6H) in *Wfs1^E864K^* homozygous and heterozygous mice, with shorter MAMs only in homozygous mice, showing, one more time, a dose dependent effect of the *Wfs1^E864K^* protein (Figure 6F-I).

## Differential gene expression in the ON

From our histological and functional data, we hypothesize that the axonal optic nerve damages precede those observed in the cell bodies of the RGCs, pointing toward a retrograde degeneration pathway. Therefore, to investigate the pathophysiological mechanisms occurring in the optic nerve, and that could explain the mitochondrial dysfunction, we performed an unbiased RNAseq analysis on isolated ONs and compared the transcriptomes of *Wfs1^WT^* and *Wfs1^E864K^* mice at P30.

Principal component analysis demonstrates a clear segregation between wild-type littermates (grey dots), and *Wfs1^E864K^* (pink dots) samples along the first principal component (PC1), with a significant total variance of 77%. This separation indicates that genotype, *i.e* wild-type littermates *versus Wfs1^E8^*^64^ is the primary source of transcriptional variation. Wild-type littermates samples cluster tightly, reflecting consistent transcriptomic profiles, whereas *Wfs1^E86K4^* samples show greater dispersion, suggesting increased biological or transcriptional heterogeneity within this group (Supplemental Figure 7). Among the 50 differentially expressed genes (DEGs), an equal distribution of up- and down-regulated genes was observed as 26 genes were upregulated in the *Wfs1^E864K^* optic nerves (Figure 7A, Supplemental Table 2), and 24 downregulated (Figure 8A, Supplemental Table 3).

**Figure 7.**
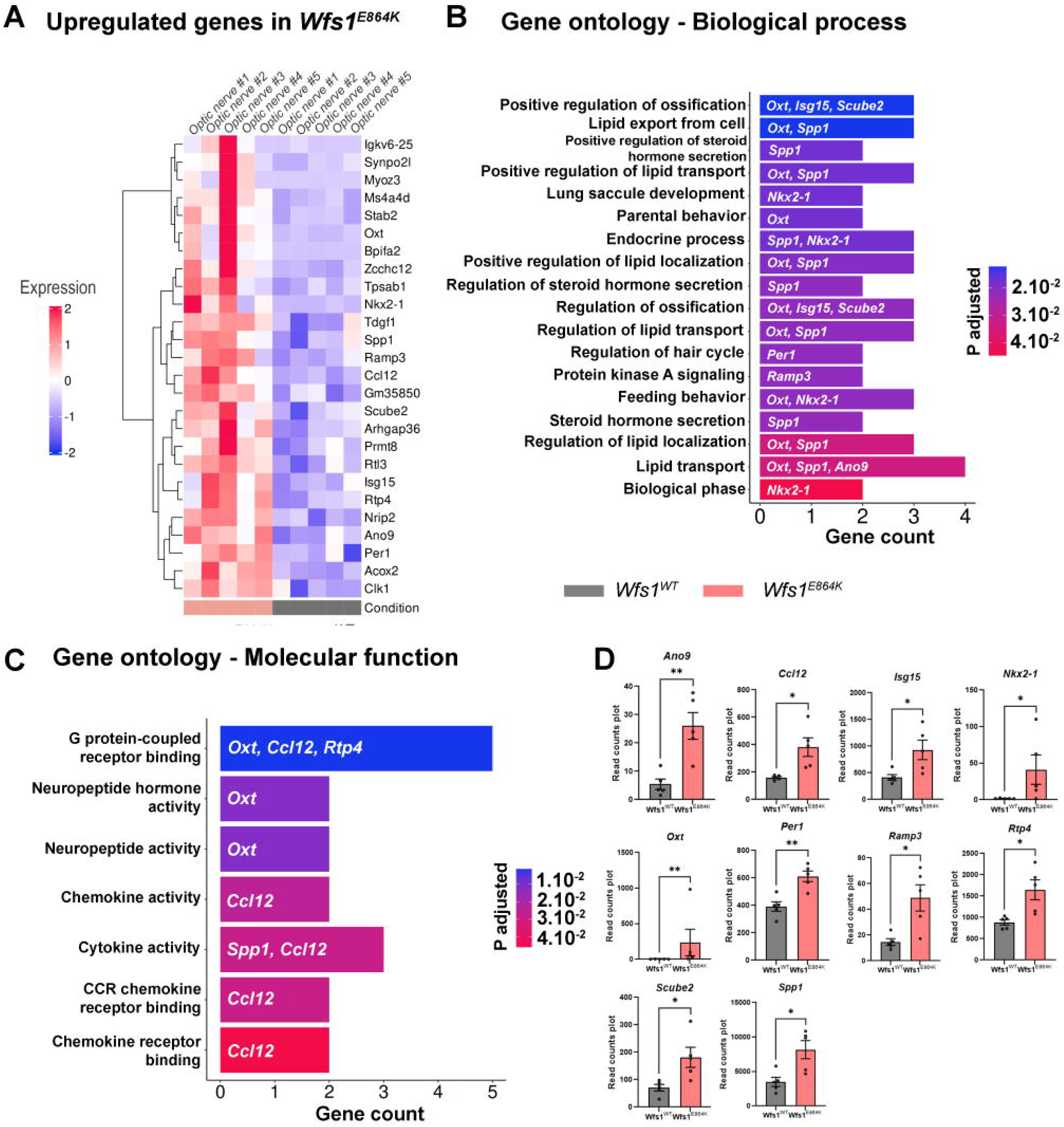
Analysis of upregulated genes in *Wfs1^E864K^*. (A) Heatmap showing the variability of expression for each sample (n=5/condition) for the 26 upregulated genes in *Wfs1^E864K^* versus *Wfs1^WT^* optic nerves. (B) Gene Ontology enrichment analysis for biological process terms of upregulated differentially expressed genes in *Wfs1^E864K^* optic nerves. The bar plot represents significantly enriched GO terms among genes upregulated in *Wfs1^E864K^* versus *Wfs1^WT^*. X-axis indicates the number of genes per term, with color representing adjusted *P* value. Representative contributing genes are listed in bar plot for each term. (C) Gene Ontology enrichment analysis for Molecular function terms of upregulated differentially expressed genes in *Wfs^E864K^* optic nerves. The bar plot represents significantly enriched GO terms among genes upregulated in *Wfs1^E864K^* versus *Wfs1^WT^*. X-axis indicates the number of genes per term, with color representing adjusted *P* value. Representative contributing genes are listed in bar plot for each term. (D) RNA-seq-based read count plot gene expression analysis of upregulated genes of interest in *Wfs1^E864K^* and *Wfs1^WT^* optic nerves. Data are presented as mean ± SEM. Unpaired t tests or Mann–Whitney tests were performed between the *Wfs1^E864K^* and *Wfs1^WT^* condition. \**p* < 0.05, \*\**p* < 0.01.

**Figure 8.**
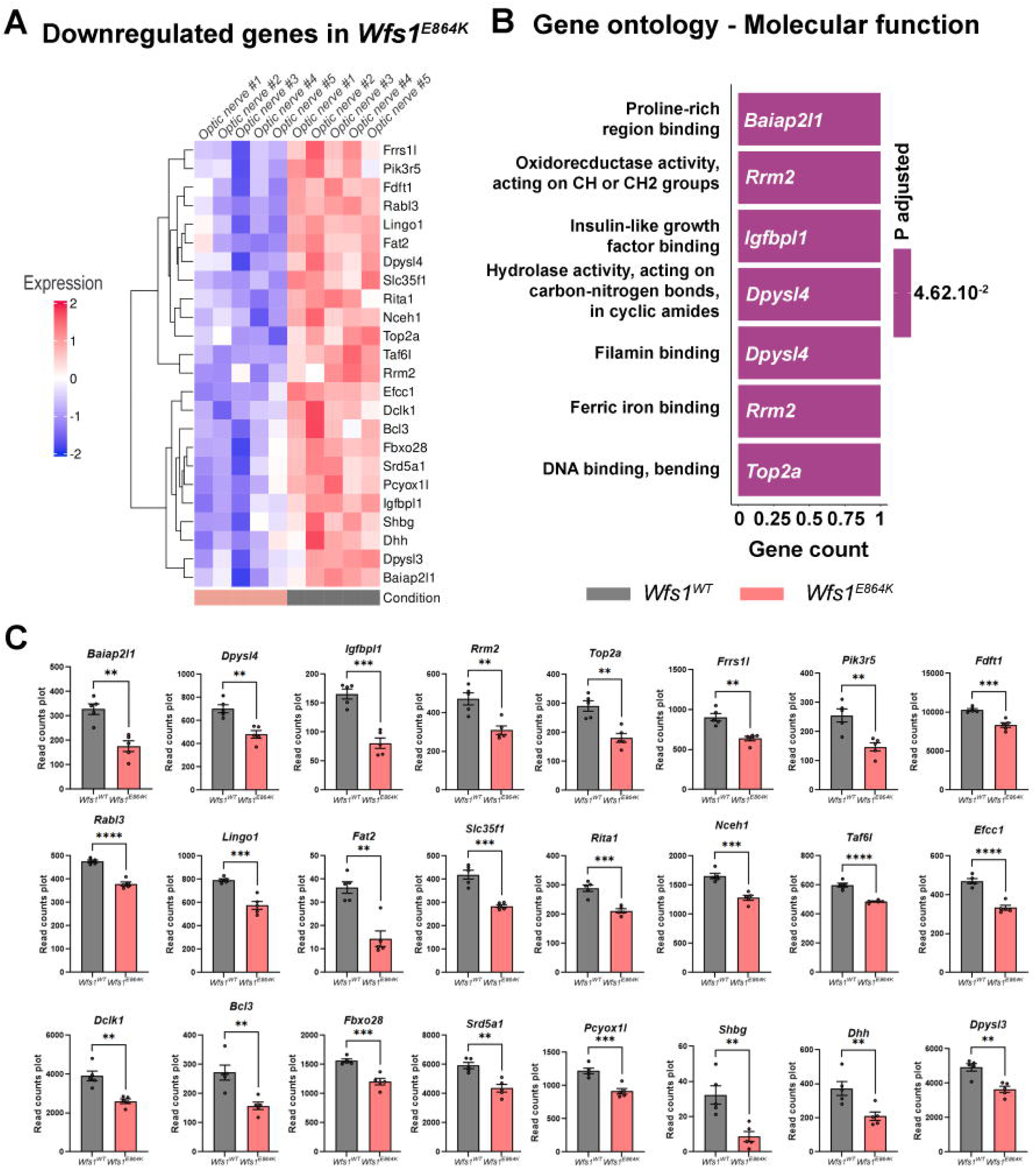
Analysis of downregulated genes in *Wfs1^E864K^*. (A) Heatmap showing the variability of expression for each sample (n=5/condition) for the 24 upregulated genes in *Wfs1^E864K^* versus *Wfs1^WT^* optic nerves. (B) Gene Ontology enrichment analysis for Molecular function terms of downregulated differentially expressed genes in *Wfs1^E864K^* optic nerves. The bar plot represents significantly enriched GO terms among genes downregulated in *Wfs1^E864K^* versus *Wfs1^WT^*. X-axis indicates the number of genes per term, with color representing adjusted *P* value. Representative contributing genes are listed in bar plot for each term. (C) RNA-seq-based read count plot gene expression analysis of downregulated genes in *Wfs1^E864K^* and *Wfs1^WT^* optic nerves. Data are presented as mean ± SEM. Unpaired t tests or Mann–Whitney tests were performed between the *Wfs1^E864K^* and *Wfs1^WT^* condition. \*\**p* < 0.01, \*\*\**p* < 0.005, \*\*\*\**p* < 0.0001.

Interestingly, combining gene enrichment analyses (Gene Ontology, GO) and literature review, we put in light up-regulated genes involved in lipid metabolism (*Oxt, Spp1* and *Ano9*), glia activation response (*Ccl12, Isg15, Scube2* and *Spp1*), extracellular matrix modeling (*Oxt, Per1* and *Nkx2-1*) and metabolic stress (*Spp1, Gal* and *Oxt*) (Figure 7B-7D). Likewise, down-regulated genes are related to cytoskeleton stability (*Balap2l1, Dpysl4, Fat2, Dclk1* and *Dpysl3*), cell survival and axonal regeneration (*Bcl3, Top2a, Pik3r5, Fat2, Rmr2* and *Pclk1*), intracellular transport (*Slc35f1*), demyelination (*Dhh*) and metabolic stress *(Rmr2* and *Pik3r5*) (Figure 8B-8C).

In addition to the DEGS, four other genes caught our attention as they are related to microtubule associated protein (*Map1b* and *Map6*) and autophagy/mitophagy dysfunction (*Tfeb* and *Lamp1*), two molecular functions of particular interest in the context of vision loss and WFS1 related pathologies. Even though their p adjusted value is over 0.05, their p value is statistically significant, with a decrease of their expression in the mutant mice compared to their wild-type controls (Supplemental table 4).

## Discussion

In the context of WLS, a rare dominant autosomal disease associated with hearing loss, OA and DM, 87% of the affected individuals develop OA, the most frequent and common ophthalmological feature in this disorder, deemed critical for the diagnosis and the follow-up of the disease progression ^9,24,28^.

In this study, we confirmed that *WFS1^E864K^* variant leads to optic disk pallor in two unrelated affected individuals as well as OPL lamination and decrease in RFNL, in line with ophthalmological observations reported by other authors ^6,8,28^. To note, case NZ01 has bilateral color vision defects, a visual deficit commonly reported with other WLS disease causing variants ^9^. Remarkedly, contrary to previous cases ^6,8^, despite clinical alterations, none of the patients had visual complaints and no severe characteristic symptoms of optic atrophy, such as visual field loss or loss of central visual acuity, were reported. This may be attributed to the young age of the affected individuals of our study. Indeed, in the study by Kobayashi et al, the 2 individuals with optic atrophy were aged 29 years and older at the time of the study, while the 2 individuals without OA were under 10 years old ^42^. Altogether, this is strengthening the importance of visual exploration in the presence of *WFS1^E864K^* variant or other WLS variant. A follow-up of the progression of the disease is critical as visual function loss might get unnoticed in the youngest but need appropriate diagnosis and care.

In a mouse model harboring the same pathological variant, the *Wfs1^E864K^* transgenic line, severe retina dysfunctions were measured using ERG. A decrease in pSTR, phNR, scotopic and photopic b-wave, and OPs amplitudes unveiled alteration of the RGCs, bipolar and amacrine cells. These electrophysiological data were corroborated by a thorough analysis of the retina layers and cell types, using various histological approaches. Thus, a progressive thinning of the inner retina and outer plexiform layers was associated with a loss of RGCs, highlighting the RGCs as the initially affected cells. RGCs have long axons that extend into the brain and form the optic nerve. Preceding retina damages, we observed, in the ON, a progressive demyelination of the axons, death of nerve fibers, with no overt loss of glial cells. Knowing the interplay between WFS1 and mitochondria, we explored their function and ultrastructure and demonstrated that mitochondria bioenergetics and structure were altered in the ON, while only a decrease of the MAMs was reported within the soma of the RGCs. These defective mitochondria could be the result of various underlying mechanisms including deregulated mitochondria transport, abnormal lipid metabolism or transport, autophagy/mitophagy processes as unveiled by a transcriptomic analysis of the ON of both *Wfs1^E864K^* homozygous mice and their wild-type counterparts.

Interestingly, in the mouse, the first signs of retinal histological damage were a thinning of the OPL as early as P30 in homozygous mutant mice, and at a later time point, P90, in the heterozygous, with a progression in time in both cases. No sign of outer retina (OS and ONL) alteration was observed during the course of our study. The inner retinal-limited alteration induced by the *Wfs1^E864K^* variant is consistent with the electrophysiological dysfunction found in RGCs, amacrine and bipolar cells. Some studies have established a positive correlation between RGCs loss and RFNL thickness in optic neuropathies such as WLS ^28,57^. The significant reduction of RGC cell density measured in a Wfs1^E864K^ dose dependent manner in our mouse model is therefore in line with the RFNL thinning of affected individuals. In addition, in the homozygous mice, a decrease of the synaptophysin immunoreactivity, which could be used to explore the plexiform layers ^58^, was measured, coinciding with the lamination of the OPL reported in human. Thus, our mouse model recapitulates the OPL alteration and RFNL decrease measured in the 2 affected individuals of this study, but also the thinning of the inner retina layer reported by others in WLS case reports (de Muijnck et al., 2024; Majander et al., 2016, Kiraly et al., 2024; O’Bryhim et al., 2022). In addition, the optic atrophy is the most characterized feature of WLS pathology ^6–8,28^, highlighted in our study with the optic disk pallor observed in both affected individuals and the thinner ON fibers in case report NZ01. In our model, the decreased number of RGCs and the thinning of the FL suggested a possible optic nerve alteration. Looking thoroughly, using a combination of histological methods, we measured a loss of axons, with a particular susceptibility to the pathogenic Wfs1^E864K^ protein in the bigger axons, preceded by myelin damage, in both mutant mice. Altogether, these phenotypic similarities, between human and mouse, support the relevance of our murine model for the study of *WFS1^E864K^* related WLS but also of WLS associated with other variants, and provide us with a model of choice to investigate the natural history of the disease.

However, contrary to what was reported for the characterization of hearing loss in this model ^41^, the heterozygous mutant mice, *Wfs1^E864K/WT,^* showed signs of a defective visual pathway as early as P30 for some parameters explored. Overall, visual impairments are more moderate than in homozygotes, presenting impairments similar to those found in humans. The progression of these alterations has also a chronology closer to that of the affected individuals, providing us with a more accurate experimental model. Thus, the transgenic mouse line offers the advantageous opportunity to explore two developmental scenarios of WLS. The heterozygous mice are a pertinent model to follow the progression of the disease in a comparable way to the one in humans while the homozygous represent an advanced, accelerated state of the disease.

Looking for the pathophysiological mechanisms responsible for the cellular alterations, in WS experimental models, reactive gliosis and inflammation have been reported, thought to be, initially, the leading cause underlying optic atrophy in the *Wfs1^exon8del^* mouse ^36,37^. Although the role of oligodendrocytes in the OA was eventually found to be limited (Ahuja et al., 2024), in our model, proinflammatory microglia cell population was increased in the RGCs of the homozygous and heterozygous mutant mice, compared to their wild-type littermates. In the ON, an increase of the astrocyte density was detected with no alteration of the oligodendrocytes nor microglia cell populations. Similar to what was observed in the *Wfs1^exon8del^* mouse model of WS, these glia cell variations could be found in response to retina and ON damages ^59^, but not as primary causes.

In the context of WS, accumulating evidence, from our lab and others, showed the role of WFS1 in the mitochondria homeostasis ^16,38,40,53,54,60^, as well as the MAMs. WS went from being qualified a “mitochondrial disease” to a more precise denomination: “MAMopathy” ^19^. However, WS is associated with few to no WFS1 protein while in WLS, an abnormal WFS1 protein is synthesized, suggesting potentially different physiological consequences. We put in light that hearing loss, in our *Wfs1^E864K^* model, was most likely stemming from a mislocalization of WFS1 binding partner, the Na^+^/K^+^ATPase β1 subunit, a key protein for the maintenance of the endocochlear potential, leading to a defective cation transfer in the stria vascularis and its collapse ^41^. Although hearing loss seems to be the consequence of a non-mitochondrial mechanism, confirming the hypothesis of a tissue specific role for WFS1 ^41^, we showed in hippocampus and cortex cultured neurons from *Wfs1^E864K^* mouse, as well as in affected individuals’ fibroblasts, a decrease in mitochondria bioenergetics, a deregulation of the mitochondrial quality system, an alteration of the autophagic flux, associated in the murine cells with a decrease of MAM number ^12^. In addition, previous authors have suggested an association between axonal mitochondria and axon health and myelination ^61,62^, mitochondria distribution in the optic nerve being associated with the state of myelination ^63^. Therefore, in this study, using SeaHorse analyzer on ON explants and TEM analysis, we explored mitochondria status and uncovered a decrease of mitochondria number associated with a severe axonal mitochondria bioenergetics alteration as well as poorly structured mitochondria, in both homozygous and heterozygous mutant mice. We also investigated the mitochondria in the RGC soma, as these cells were strongly impacted by the Wfs1^E864K^ protein, with however more limited alterations. Although, the number as well as the integrity of the mitochondria were well preserved, MAMs disruption was observed with a decrease of their number and length in *Wfs1^E864K^* homozygous mice, and only a decrease of their number in *Wfs1^E864K/WT^* mice. These results strengthen our hypothesis that axonal mitochondria are the main players in *Wfs1^E864K^* -induced OA. Markedly, clinicians have recently suggested that, contrary to what was seen in other optic neuropathies, the optic nerve atrophy was preceding any retinal damages in both Wolfram and Wolfram-like syndromes ^28,57^. Although it is difficult to say for our two affected individuals, the study of our experimental model allows us to draw the same conclusion. Indeed, the optic nerve damages were first observed when no histological changes were found in the retina and the divergence between the structure of ON and RGC mitochondria seems to support this fact. Interestingly, exploration of optic atrophy in the *Wfs1^exon8del^*, a mouse model for WS, also concluded that optic nerve damage predominates over RGC damage ^36^.

Therefore, assuming that defects, including mitochondrial ones, first appear in the ON, we performed transcriptomics analysis between *Wfs1^E864K^* mice and their wild-type counterparts, at an early time point, P30. The *Wfs1^E864K^* variant led to an up regulation of genes involved in glia activation response (*Ccl12, Isg15, Scube2* and *Spp1*), with no changes in microglia proliferation and proinflammatory related genes (*Edn2, Itgam, Cd68, C4b, Csf1r*), consistent with the astrogliosis detected in the ON (Supplemental Figure 5C), without changes in microglia (Supplemental Figure 5E). The differential expression of genes implicated in the cell survival and axonal regenerated factors also supports the axonal loss seen at advanced stages while the lower expression of *Dhh*, a gene involved in the myelination maintenance, as well as the increase of lipid metabolism genes expression could be linked to the myelin disorganization and damage ^64^.

Surprisingly, our analysis did not highlight any variations in ER stress and Ca^2+^ homeostasis pathways, even though the role of WFS1 in these specific molecular pathways is commonly accepted ^15,17,53^. Nonetheless, *Lamp1* and *Tfeb* genes, which have been implicated in autophagy and mitophagy processes respectively ^65–68^, were found down-regulated in the *Wfs1^E864K^* mice. This deregulation is consistent with the data presented in cultured neuronal cells from this mouse line and fibroblasts from *WFS1^E864K^* carrier, where both autophagy and mitophagy processes were altered ^12^. On the other hand, genes related to lipid metabolism (*Oxt, Spp1* and *Ano9*), tissue morphogenesis and extracellular matrix remodeling (*Oxt, Per1* and *Nkx2-1*) were found up-regulated while several genes, *Balap2l1, Dpysl4, Bcl3, Top2a, Slc35f1, Map1b,* among others, linked to cytoskeleton stability, intracellular transport and microtubes impairment were down-regulated. More particularly, *Map1b* gene, coding for the microtubule-associated protein 1B (MAP1 B), which is known to be involved in cytoskeletal rearrangements by regulating actin and microtubule dynamics, has recently been suggested to be a cause of impaired axonal transport of mitochondria, in the pathophysiological context of spinal muscular atrophy ^69^. Taken together with the bioenergetics and structural deficits observed in the mitochondria of the ON, it is therefore tempting to speculate that mitochondrial dysfunction might be the result of intracellular transport defects of the mitochondria from the soma to the axons, associated with a defective quality control and mitophagy process, leading to a decrease in their number in the axons as well as a defective and damaged mitochondrial population, even though additional studies would be necessary to confirm our hypothesis.

In conclusion, *Wfs1^E864K^* variant induced early OA in mice, with RGCs loss and OPL thinning, recapitulating the visual features of *WFS1^E864K^* affected individuals, and more broadly WLS patients. This experimental model reproduces the progression of the disease in a similar chronology to WLS pathology opening two possible scenarios of damage to explore new therapeutics or mechanisms, using either the homozygous *Wfs1^E864K^* mutant mice for an advanced and severely impaired model or the heterozygous *Wfs1^E864K/WT^* mice for an early stage of the disorder. Our experimental model shed light on the implications of axonal mitochondria impairment as a main player in the pathophysiology of WLS, opening novel therapeutics perspectives in the WFS1 related vision loss.

## Methods

### Sex as a biological variable

Our study examined male and female animals, and similar findings are reported for both sexes.

### Human clinical assessment

Patient NZ01 underwent ophthalmologic evaluation, including Snellen visual acuity, Ishihara colour testing, cycloplegic refraction, wide field photography and fundus autofluorescence, (Optos plc, Dumferline, UK), and spectral domain Ocular coherence tomography (HRA+OCT Heidelberg Spectralis). Patient NL01 underwent visual acuity measurement using Snellen charts. Visual fields were assessed with Octopus automated perimetry. OCT was conducted with Canon Xephilio OCT-A1 (Canon Inc., Kanagawa, Japan). Color vision was assessed with HRR Pseudoisochromatic Plates. Pattern-reversal VEPs were recorded according to the ISCEV (International Society for Clinical Electrophysiology of Vision) standard with the Espion E3 system (Diagnosys LLC, Cambridge, UK).

### Mouse breeding and experimental design

The *Wfs1^E864K^* mice were generated as previously described ^41^. Animal care and experimentation were authorized by the national ethic committee (Paris, France) and carried out in strict adherence to European Union Directive 2010/63 and ARRIVE guidelines. The mice were housed in acrylic transparent cages with ad libitum food and water under 12:12 light/dark cycle and temperature-controlled conditions. Experiments were conducted with *Wsf1^WT^*, *Wfs1^E864K/WT^* and *Wfs1^E864K^* mice at the same time of the day during the light cycle period. Genotyping and sequencing were done by routine PCR as described previously in ^41^. The ERG recording and histology procedures were performed at post-natal day (P)30, P60 and P90 while Seahorse and RNAseq studies were done at P30 only.

### ERG recording and analysis

After dark adaptation, the mice were anesthetized with 40 mg/kg ketamine and 1 mg/kg xylazine under dim red illumination by intraperitoneal injections. Eyes were topical treated with 5 % Neosynephrine (Europhta, Monaco), Mydriaticum 2 mg/0,4 ml (Thea pharma, Clermont-Ferrand, France) and Cebesine 0,4% (Chauvin, Montpellier, France) before recording. The ERG, developed by VisioSystem (SIEM biomedical, Nimes, France) was equipped with a 400 mm Ganzfeld and white-light-emitting-diode stimulator, covered by faraday hood to avoid electrical noise. To record retinal function, wet cotton electrodes attached to recording electrodes were used (ECLIPS-AR, SIEM biomedical, Nimes, France), both reference electrodes were inserted under the skin behind the ears, and a grounding electrode was attached to the tail (PEA1S, SIEM biomedical, Nimes, France). Under scotopic conditions, the mice were exposed to 5 flashes of 14 mcd.s/m^2^ intensity for pSTR traces or 3.2 cd. s/m^2^ intensity for a-wave and b-wave traces, 3 flashes of 1.2 cd. s/m^2^. For photopic analysis, after 5 minutes of 30 cd.s/m^2^ light-adaptation, the mice were exposed to 10 flashes of 3.2 cd. s/m^2^ for photopic b-wave and phNR traces. Ultimately, the mice were place in a cage on a heat plate at 37℃ with topical Ocry-gel (TVM lab, Pont-du-chateau, France) to ensure animal health and recovery.

VisioSystem software was used for ERG recording and data acquisition. The pSTR parameter was obtained by measure the highest maximum positive peak at 14 mcd.s/m^2^. The a-wave was measured as the difference in the amplitude from the baseline to the maximum peak of the negative deflection while the b-wave was measured as the amplitude difference between the negative peak to the highest positive peak consequently at 3.2 cd. s/m^2^. The OPs waves were taken as the mean of the three maximum peaks at 1.2 cd. s/m^2^. For photopic b-wave and phNR, the maximum positive peak and the consequent negative deflection peak were measured, respectively, at 3.2 cd. s/m^2^. For statistical comparison, the mean values from both eyes of 6 to 10 mice were averaged.

### Histology procedures

Animals were deeply anesthetized using ketamine and xylazine as previously mentioned and intracardially perfused with warm NaCl 0.9% solution followed by fixation with paraformaldehyde solution (Antigenefix, Cat. P0016, DiaPath). Eyes were enucleated, the cornea were removed, and eyes were post-fixed overnight at 4°C with paraformaldehyde 4%, pH = 7,4 (GF720170-1010, Delta microscopies). Additionally, ONs were obtained from the optic nerve head to optic chiasm and post-fixed overnight at 4°C in the fixative solution described above. The samples were dehydrated, cleared in isopropanol and, after removal of the crystalline for the retinas, embedded in paraffin. Transversal section from retina and distal ON, 8 μm and 7 μm respectively, were obtained using a microtome Leica RM2145.

#### Hematoxylin and Eosin staining

Retinal sections were stained by H-E for morphometry analysis. Retina sections were dewaxed, rehydrated and stained with Mayer hematoxylin (cat. 75290, Sigma Aldrich) solution for 10 minutes. After washing in tap water and in 0.05% Li_2_CO_3_ solution (ref. PRO-25004, Prolabo), Eosin (cat. EE0190, Bio basic) 1% solution was used for counterstaining for 30 seconds. Finally, sections were dehydrated and mounted (cat. 00811-EX, PERTEX®, HistoLab) for light microscopic acquisition.

Light microscopic images were captured with an up-right microscope (Leica DM 2500) equipped with a DFC495 camera using a 40 X objective. Using Image J software, the thickness of the total retinal and each layer (OS/IS, ONL, OPL, INL and IPL) was measured from two opposite areas of the retina at 400 mm from the optic nerve head, and the average of both values was computed. The mean value from 6 - 10 eyes were averaged for statistical comparison. The mean value from 3 images from 5 – 10 eyes per genotype were computed.

#### Luxol Fast Blue staining

Transversal ON sections were stained with LFB for myelin content studies. ON sections were dewaxed, hydrated and incubated overnight in 0.1% LFB solution (cat. 212170250, Termo Fisher Scientific) at 60℃. Sections were washed with ethanol 70% for 10 seconds to rinse off the excess of luxol fast blue. After, differentiation was carried out with 0.05% Li_2_CO_3_ followed by ethanol 70% for 30 seconds each. Sections were dehydrated and mounted for light microscopic acquisition. Light microscopic acquisition was performed using an upright microscope Leica DM 2500 equipped with a DFC495 camera.

### Immunofluorescence procedures

#### Immunofluorescence on sections

Immunofluorescence studies were performed in ON and retina paraffin sections. After dewaxing, antigen retrieval was done using citric acid buffer (citric acid monohydrated, Cat. 27490, Fluka) 0,2 p/v % with tween 20 (Cat. 2001-C, Euromedex), pH = 6 at 90℃, for 30 minutes. After washing with PBS, the sections were blocked with 10% horse serum in PBS – Triton 100 0.2% for 2 hours, followed by an overnight incubation with primary antibodies at 4℃ in blocking solution. After several washes with PBS, sections were incubated with secondary antibody in blocking solution for 2 hours at room temperature: donkey anti-rabbit Alexa 567 and donkey anti-mouse Alexa 488 (1:400, A10042 and A21202, Termo Fisher Scientific). Subsequently, nuclei were stained using DAPI (D9542-50MG, Sigma Aldrich) for 10 minutes. Finally, sections were washed with PBS and mounted (Dako fluorescent mounting medium, Agilent Technologies, France) for confocal acquisition. The following primary antibodies were used: Cleaved Caspase-3 (1:300, 9661S, Cell signaling), Rhodopsin (1:500, ab98887, Abcam), Cone-Arrestin (1:500, ab15282, Millipore), PKC-1a (1:100, MA1-157, Invitrogen), GFAP (1:500, 123895, Cell signaling), Synaptophysin (1:500, ab14692, Abcam), Oligo-2 (1:200, MABN50, Sigma Aldrich), IBA-1 (1:500, 019-19741, Wako) and TUJ1 (1:500, 801201, Biolegend). All acquisitions were performed with a confocal microscope LSM880 ZEISS (MRI-DBS, Montpellier, France).

#### Retina flat mount immunofluorescence

The eyes were post-fixed with PFA 4% for 60 minutes at room temperature following the enucleation and removal of cornea and crystalline. Dissection of the eyes were performed, and retinas were detached and flat mounted without the vitreous. Subsequently, retinas were permeabilized by freezing at −80℃ for 30 minutes. After 2 hours of blocking with 5% horse serum in PBS-Triton 100 0.5%, incubation with the following primary antibodies was performed for 48 hours, at 4°C, in blocking solution: Brn3a (1:500, sc-8429, Santa Cruz), IBA-1 (1:500, 019-19741, Wako) and CD68 (1:300, ab318303, Abcam). After washing with PBS, retinas were incubated with aforementioned secondary antibody for 2 hours at room temperature. Nuclei were stained with DAPI and after several washes with PBS, retinas were mounted using mounting medium (Dako fluorescent mounting medium, Agilent Technologies, France) for confocal acquisition.

#### Immunofluorescence analysis using Python

The analysis of immunoreactive positive area (PKC, GFAP, Synaptophysin, Rhodopsin, Cone-Arrestin and TUJ1), IBA1 and CD68 colocalization and identification of number of cells (Oligo2, IBA-1 and Brn3a) was performed in retina and ON transversal confocal images captured at 20 x by Python script available online (GitHub). For analysis of immunoreactivity positive area in retina and ON sections, brightness and contrast of the images were adjusted using Adobe Photoshop. For IBA1&CD68 colocalization index the number of positive pixels was quantified on identified cells. For quantification of the number of cells, the background was reduced and the signal enhanced by the Python script followed by contour identification. Images were analyzed under equal binarization threshold and the values from 5 – 7 eyes per genotype were averaged.

### Ultrastructure analysis by electron transmission microscopy

#### Sample preparation

After animal perfusion has previously described, enucleated eyes (without the cornea and crystalline) and ONs were immersed in a solution of 2.5% paraformaldehyde + 2.5% glutaraldehyde in cacodylate buffer (Cat. GF210174-1010, Delta microscopies) overnight at 4°C. They were then rinsed in cacodylate buffer and post-fixed in a 0.5% osmic acid + 0.8% potassium Hexacyanoferrate trihydrate in water for 2 h at dark and room temperature. After two washes in water, the cells were dehydrated in a graded series of ethanol solutions (30-100%). The samples were embedded in EmBed 812 using an Automated Microwave Tissue Processor for Electronic Microscopy, Leica EM AMW. Thin sections (70 nm; Leica-Reichert Ultracut E) were collected at different levels of each block. These sections were counterstained with uranyl acetate 1.5% in 70% Ethanol and lead citrate and observed using a Tecnai F20 transmission electron microscope at 120KV in the Institut des Neurosciences de Montpellier: Electronic Microscopy facilities, INSERM U 1298, Université Montpellier, Montpellier France.

#### Axon classification and quantification

The quantification of the number of axons was performed in ultrastructural images taken at 3000x by TEM microscope using the following python script available online (GitHub). The analysis was done by binarization, after identifying the contours of each axon, the total number of axons and the number of axons of three different sizes were quantified: small axons (axon diameter between 0,3 - 0.8 mm), medium axons (axon diameter: 0.8 – 1.4 mm) and larger axons (1.4 – 2.2 mm). The selected area and the binarization threshold were chosen in order to have the best representation of axons with the lowest bias. The mean value of 3 images from 5 – 7 eyes per genotype were computed.

#### Myelin disorganization and g-ratio analysis

Quantification of the disorganization of myelin and g-ratio were performed in ultrastructural images taken at 10000x and 25000x by TEM microscope. The number of elongated and uncompacted myelin was identified and count as an event of disorganization. The number of disorganized events per image was analyzed and the mean value from 3 images from 5 – 7 ONs per genotype were computed. For g-ratio quantification, two to four axons of different sizes were selected per image and using ImageJ software, three thickness measurements were average per axon. The mean value of 3 images from 5 – 7 ONs per genotype were computed.

#### Mitochondria ultrastructure in the optic nerve

Using Tecnai F20 transmission electron microscope, images from ON transversal sections were taken at 10,000 x and 25,500 x. The number of myelin disorganization events per image, the g-ratio (length of the myelin sheath) and mitochondria ultrastructural analysis were done using Image J software. The number of mitochondria per image, their perimeter and elongation factor were computed. The differences in electron density were used to reveal the presence of mitochondria cristae, images (25,500 x) were binarized under specific threshold that represented the area occupy by cristae, then the percentage of cristae area was normalized to the mitochondria area, and cristae complexity score was computed from 1 to 5 considering 1 point of score every 20% area occupy by cristae. The mean value from 5 images from 6 eyes per genotype was calculated for myelin disorganization and g-ratio at 10,000 x and 25,500 x, respectively. The number of mitochondria was counted 5 images (10,000 x) from 6 eyes per genotype, while mitochondria 50 – 70 from 4 images (25,500 x) from 5 to 7 ONs per genotype, were analyzed and values were averaged for the other parameters.

#### MAMs ultrastructure in RGCs

Images from retina transversal sections were captured at 25,500 x by Tecnai F20 transmission electron microscope. The mitochondria analysis was done as previously described. Additionally, the mitochondria and ER from the RGCs were identified and the number and length of MAMs (ER-mitochondria distance between 10 μm and 30 μm) were analyzed with Image J, in 4 images from 5 eyes per genotype between 75 to 100 mitochondria were quantified.

### RNA-seq procedures

#### RNA extraction and RNA-seq libraries from ON mice samples

For whole transcriptome analysis, total RNA was extracted from optic nerves *Wfs1^WT^* (n=5) or *Wfs1^E864K^* (n=5) after tissue lysis using the NucleoSpin RNA XS kit (Macherey–Nagel, ref.740902.250) according to the manufacturer’s protocol. Total RNA was quantified, and its quality was determined using a BioAnalyser 2100 with the RNA 6000 PICO Kit (Agilent Technologies, Leuven, Belgium). RNA samples with a RIN >7.8 were selected for RNA-seq analysis. mRNA libraries were prepared following the manufacturer’s recommendations (SmartSeq V4 Ultra Low Input RNA kit from TAKARA). The final library preparation of pooled samples was sequenced on a Nextseq 2000 ILLUMINA device with a P2-200 cycle cartridge (2 × 400 million 100-base reads) corresponding to 2 × 40 million reads per sample after demultiplexing.

#### RNA-Seq data analysis and bioinformatic analysis of biological processes and pathways

Fastq files obtained from the sequencing were aligned using STAR (v2.7.9a) against the Mouse reference genome from Ensembl v110 (generated 2023/08/29), with the option “quantMode GeneCounts” to extract the raw counts for each gene, and all count files were concatenated into a single file. A sample file was created with the sample data, including the conditions (*Wfs1^WT^*, *Wfs1^E864K^*) and number of replicates. The count file and sample file were loaded into our inhouse R Shiny application EYE DV seq (Plateforme Phenotypage Cellulaire et Tissulaire, Institut de la Vision, Paris, France). We added 1 to all counts in the count file to avoid any 0-read count errors. We then removed the genes with a total count of < 10 for all samples. Finally, DESeq2 (v1.40.2) analysis was performed comparing the groups defined in the ‘condition’ column, with ‘*Wfs1^WT^*’ as the ‘control condition’ and ‘*Wfs1^E864K^*’ as the ‘treatment condition’. The results were then filtered for significant genes using the *p*-values (< 0.05, base mean minimum 10, none log_2_ fold change exclusion) in order to have an optimal scan of the sequencing, allowing us to obtain of 50 differentially expressed genes (DEG). An enrichment analysis was performed to determine the involve pathways using the GO database (using the Bioconductor packages ‘org.Rn.eg.db’ (v3.17.0) and ‘clusterProfiler’ (v 4.8.1)). Enrichment analyses were generated based on Gene Ontology database focusing on Biological Process and Molecular Function of DEG (exclusion of log_2_ fold change between – 0.5 and 0).

### Determination of oxygen consumption rate (OCR) in optic nerve

OCR were evaluated in isolated ONs using Seahorse XFe24 analyzer (Agilent Seahorse XF) as described in a previous report ^56^. Each Seahorse XF24 Sensor cartridge plate was incubated with calibrated buffer (cat. 100840-000, Agilent Seahorse XF) at 37℃ CO_2_-free incubator one day before the experiment. Animals were euthanized by cervical dislocation, and the ONs were taken as mentioned previously and rapidly incubated in DMEM (Cat. 102353-100, Agilent Seahorse XF) supplemented with 0.5 mM sodium pyruvate (Cat. 103578-100, Agilent Seahorse XF), 2 mM glucose (103577-100, Agilent Seahorse XF) and 4 mM glutamine (Cat. 103579-100, Agilent Seahorse XF) at 37℃. Both ONs from one animal were dissected into small pieces of ∼ 1 mm using a surgical blade under a light surgical microscope Leica MZ 6. Between 8 – 12 ON pieces were secured to the mesh insert of a Seahorse XF24 Islet Capture Microplate (Cat.101122-100, Agilent Seahorse XF) contained warmed supplemented DMEM. To evaluate the mitochondrial function in the ON with or without (or reduced) mitochondria glial contribution, oxygen consumption rates were measured without or with barium fluorocitrate administration (Cat. F9634-100MG, Sigma Aldrich), an inhibitor of the aconitase from the glial cells ^55,56,70^. A 2,5 mM Flc solution was performed according to previous reports ^56^. Ports were loaded with the following compounds: Port A: DMEM or fluorocitrate barium (2,5 mM); Port B: Oligomycin (100 mg/ml; Cat. 103015-100, Agilent Seahorse XF); Port C: FCCP (44 mM; Cat. 103015-100, Agilent Seahorse XF) and sodium pyruvate (100 mM); Port D: Rotenone/antimycin A (130 mM; Cat. 103015-100, Agilent Seahorse XF). Following each Seahorse XF24 Sensor Cartridge being calibrated, the experiment was run in supplemented DMEM to measure OCR ON by following sequential components: A) DMEM; B) Oligomycin; C) FCCP; D) Rotenone/antimycin. To measure mitochondria OCR, (without or with less mitochondria glial contribution) predominantly in neuronal axons, the following sequential components were injected: A) Flc; B) Oligomycin; C) FCCP; D) Rotenone/antimycin.

Finally, the ONs recovered from the Seahorse plates and homogenized with RIPA buffer (Tris 50 mM pH = 7,5, 150 mM NaCl, 1mM EDTA, 3 mM EGTA, 1 Mm DTT, SDS 0,1 % p/v, sodium deoxycholate 0,1 %) with protease (Protease inhibitor cocktail; Cat. 11836153001, Roche) and phosphatase inhibitors (PhosSTOP; Cat. 4906845001, Roche). Homogenates were then centrifuged at 0.9 rfc for 10 minutes at 4°C and the supernatants were taken to measure the total protein concentration with a BCA assay (23227, Termo Fischer Scientific).

To evaluate OCR without or with fluorocitrate inhibitor, the mitochondria basal respiration was measured taking the mean of three last points post-injection of DMEM or Flc by port A. The proton-leak, ATP production, maximal capacity and non-mitochondrial respiration rates were measured by taking the mean of four points at each step. Finally, all data were normalized to the total protein concentration obtained with the BCA assay. The mean value of both ONs from 6 – 8 eyes per genotype were computed.

### Statistical analysis

Data are expressed as mean ± standard error to the mean (SEM), if not stated otherwise, in graph plotted using Matplotlib package from Python. Statistical analysis was performed using scikit-learnt, SciPy and Statsmodels packages from Python. Statistical significance between groups was determined by one-way ANOVA followed by post hoc Tukey’s test or Kruskal Wallis test followed by Dunn test. For the analysis of g-ratio and axon diameter, Spearman test was performed to analyze association in each group and ANCOVA analysis was done for comparison between groups. For RNAseq data analysis, statistical analysis was performed using Graph-Pad Prism 10. Unpaired t-tests or Mann–Whitney tests were performed for unpaired comparisons of RNA-seq data analysis. The levels of statistical significance considered were: * *p* < 0.05, ** *p* < 0.0, *** *p* < 0.001 and **** *p* < 0.0001.

### Study approval in human

All clinical investigation has been conducted according to Declaration of Helsinki principle. This study was assessed and approved by the medical ethical testing committee of Amsterdam UMC, location AMC, as well as with ethics approval NTX/08/12/123 (Ministry of Health), A+4290 (Te Toka Tumai). The patients and their guardians provided written informed consent.

### Data availability

Data will be made available upon request.

## Acknowledgements

We acknowledge the imaging facility MRI, member of the national infrastructure France-BioImaging (https://ror.org/01y7vt929) supported by the French National Research Agency (ANR-24-INBS-0005 FBI BIOGEN. We thank Zebrasens and Ze-neuro from the Aquatic model platform ZEFIX”. This work benefited from equipment and services from the iGenSeq core facility (Genotyping and sequencing) at ICM. We thank Xavier Guillonneau and Frederic Blond from the Institut de la Vision for their contribution in RNA-seq data extraction and analysis.

The work was supported by the AFM-Téléthon #24197, French National Research Agency ANR-23-CE14-0087 to BD, the Bartiméus Fonds #1219277, The Netherlands, to CDM, Save Sight Society NZ to ALV.

## Author Contributions

BD conceived the project. HHD, EMR, SMP and BD designed the experiments. CDM, MVG, ALV provided patient data. HHD, KD, ER, JS, CC, SA, JD, VF contributed to acquisition, analysis, and interpretation of the data. HHD, ER and EMR wrote the manuscript, and HHD, KD, ER, CDM, MVG, ALV, JS, CC, SA, JD, VF, SP, SMP, BD revised the manuscript.

## References

1. Pallotta, M. T. et al. Wolfram syndrome, a rare neurodegenerative disease: From pathogenesis to future treatment perspectives. J. Transl. Med. 17, 1–12 (2019).

2. Urano, F. Wolfram Syndrome: Diagnosis, Management, and Treatment. Curr. Diab. Rep. 16, 1–8 (2016).

3. Barrett, T., Bundey, S. E. & Macleod, A. F. (DIDMOAD) syndrome UK nationwide study of Wolfram. Lancet 346, 1458–1463 (1995).

4. Fraser, F. C. & Gunn, T. Diabetes mellitus, diabetes insipidus, and optic atrophy. An autosomal recessive syndrome? J. Med. Genet. 14, 190–193 (1977).

5. Medlej, R. et al. Diabetes mellitus and optic atrophy: A study of Wolfram syndrome in the Lebanese population. J. Clin. Endocrinol. Metab. 89, 1656–1661 (2004).

6. Eiberg, H. et al. Autosomal dominant optic atrophy associated with hearing impairment and impaired glucose regulation caused by a missense mutation in the WFS1 gene. J. Med. Genet. 43, 435–440 (2006).

7. Rendtorff, N. D. et al. Identification of p.A684V missense mutation in the WFS1 gene as a frequent cause of autosomal dominant optic atrophy and hearing impairment. Am. J. Med. Genet. Part A 155, 1298–1313 (2011).

8. Valéro, R., Bannwarth, S., Roman, S., Paquis-Flucklinger, V. & Vialettes, B. Autosomal dominant transmission of diabetes and congenital hearing impairment secondary to a missense mutation in the WFS1 gene. Diabet. Med. 25, 657–661 (2008).

9. de Muijnck, C., Brink, J. B. te., Bergen, A. A., Boon, C. J. F. & van Genderen, M. M. Delineating Wolfram-like syndrome: A systematic review and discussion of the WFS1-associated disease spectrum. Surv. Ophthalmol. 68, 641–654 (2023).

10. Kõks, S. Genomics of Wolfram Syndrome 1 (WFS1). Biomolecules 13, 1–12 (2023).

11. Rigoli, L., Caruso, V., Salzano, G. & Lombardo, F. Wolfram Syndrome 1: From Genetics to Therapy. Int. J. Environ. Res. Public Health 19, 1–18 (2022).

12. Patergnani, S. et al. The Wolfram-like variant WFS1E864K destabilizes MAM and compromises autophagy and mitophagy in human and mice. Autophagy 20, 2055–2066 (2024).

13. Gong, Y., Xiong, L., Li, X., Su, L. & Xiao, H. A novel mutation of WFS1 gene leading to increase ER stress and cell apoptosis is associated an autosomal dominant form of Wolfram syndrome type 1. BMC Endocr. Disord. 21, 1–13 (2021).

14. Bonnet Wersinger, D., et al. Impairment of visual function and retinal ER stress activation in Wfs1-deficient mice. PLoS One 9, (2014).

15. Takei, D. et al. WFS1 protein modulates the free Ca2+ concentration in the endoplasmic reticulum. FEBS Lett. 580, 5635–5640 (2006).

16. Zatyka, M. et al. Depletion of WFS1 compromises mitochondrial function in hiPSC-derived neuronal models of Wolfram syndrome. Stem Cell Reports 18, 1090–1106 (2023).

17. Liiv, M. et al. ER calcium depletion as a key driver for impaired ER-to-mitochondria calcium transfer and mitochondrial dysfunction in Wolfram syndrome. Nat. Commun. 15, (2024).

18. Erustes, A. G. et al. Overexpression of α-synuclein inhibits mitochondrial Ca2+ trafficking between the endoplasmic reticulum and mitochondria through MAMs by altering the GRP75–IP3R interaction. J. Neurosci. Res. 99, 2932–2947 (2021).

19. Delprat, B., Maurice, T. & Delettre, C. Wolfram syndrome: MAMs’ connection? review-article. Cell Death Dis. 9, (2018).

20. Ustaoglu, M., Onder, F., Karapapak, M., Taslidere, H. & Guven, D. Ophthalmic, systemic, and genetic characteristics of patients with Wolfram syndrome. Eur. J. Ophthalmol. 30, 1099–1105 (2020).

21. Kabanovski, A., Donaldson, L. & Margolin, E. Neuro-ophthalmological manifestations of Wolfram syndrome: Case series and review of the literature. J. Neurol. Sci. 437, 120267 (2022).

22. Hoekel, J. et al. Ophthalmologic correlates of disease severity in children and adolescents with Wolfram syndrome. J. AAPOS 18, 461–465.e1 (2014).

23. Zmyslowska, A. et al. Delayed recognition of wolfram syndrome frequently misdiagnosed as type 1 diabetes with early chronic complications. Exp. Clin. Endocrinol. Diabetes 122, 35–38 (2014).

24. Kiraly, P., Reichel, F. F. & Fischer, M. D. Multimodal imaging in autosomal dominant Wolfram syndrome and long-term follow-up of laminations of the outer plexiform layer. Eye 38, 110–111 (2024).

25. Hoekel, J. et al. Visual pathway function and structure in Wolfram syndrome: Patient age, variation and progression. BMJ Open Ophthalmol. 3, 1–6 (2018).

26. Zmyslowska, A. et al. Retinal thinning as a marker of disease progression in patients with wolfram syndrome. Diabetes Care 38, e36–e37 (2015).

27. O’Bryhim, B. E. et al. Longitudinal Changes in Vision and Retinal Morphology in Wolfram Syndrome. Am. J. Ophthalmol. 243, 10–18 (2022).

28. de Muijnck, C. et al. Characteristics of autosomal dominant WFS1-associated optic neuropathy and its comparability to OPA1-associated autosomal dominant optic atrophy. Sci. Rep. 14, 1–9 (2024).

29. Majander, A. et al. Lamination of the Outer Plexiform Layer in Optic Atrophy Caused by Dominant WFS1 Mutations. Ophthalmology 123, 1624–1626 (2016).

30. Schmidt-Kastner, R. et al. Expression of the diabetes risk gene wolframin (WFS1) in the human retina. Exp. Eye Res. 89, 568–574 (2009).

31. Kawano, J., Tanizawa, Y. & Shinoda, K. Wolfram syndrome 1 (WFS1) gene expression in the normal mouse visual system. J. Comp. Neurol. 510, 1–23 (2008).

32. Yamamoto, H. et al. Wolfram syndrome 1 (WFS1) protein expression in retinal ganglion cells and optic nerve glia of the cynomolgus monkey. Exp. Eye Res. 83, 1303–1306 (2006).

33. Morikawa, S., Blacher, L., Onwumere, C. & Urano, F. Loss of Function of WFS1 Causes ER Stress-Mediated Inflammation in Pancreatic Beta-Cells. Front. Endocrinol. (Lausanne). 13, 1–12 (2022).

34. Plaas, M. et al. Wfs1-deficient rats develop primary symptoms of Wolfram syndrome: Insulin-dependent diabetes, optic nerve atrophy and medullary degeneration. Sci. Rep. 7, 1–16 (2017).

35. Cairns, G. et al. A mutant wfs1 zebrafish model of Wolfram syndrome manifesting visual dysfunction and developmental delay. Sci. Rep. 11, 1–12 (2021).

36. Rossi, G. et al. MCT1-dependent energetic failure and neuroinflammation underlie optic nerve degeneration in Wolfram syndrome mice. Elife 12, 1–25 (2023).

37. Ahuja, K. et al. A deep phenotyping study in mouse and iPSC models to understand the role of oligodendroglia in optic neuropathy in Wolfram syndrome. Acta Neuropathol. Commun. 12, (2024).

38. Crouzier, L. et al. NCS1 overexpression restored mitochondrial activity and behavioral alterations in a zebrafish model of Wolfram syndrome. Mol. Ther. Methods Clin. Dev. 27, 295–308 (2022).

39. Crouzier, L. et al. Convolamine, a tropane alkaloid extracted from Convolvulus plauricalis, is a potent sigma-1 receptor-positive modulator with cognitive and neuroprotective properties. Phyther. Res. 38, 694–712 (2024).

40. Crouzier, L. et al. Morphological, behavioral and cellular analyses revealed different phenotypes in Wolfram syndrome wfs1a and wfs1b zebrafish mutant lines. Hum. Mol. Genet. 31, 2711–2727 (2022).

41. Richard, E. M. et al. Wfs1 E864K knock-in mice illuminate the fundamental role of Wfs1 in endocochlear potential production. Cell Death Dis. 14, (2023).

42. Kobayashi, M. et al. WFS1 mutation screening in a large series of Japanese hearing loss patients: Massively parallel DNA sequencing-based analysis. PLoS One 13, 1–19 (2018).

43. Fukuoka, H., Kanda, Y., Ohta, S. & Usami, S. I. Mutations in the WFS1 gene are a frequent cause of autosomal dominant nonsyndromic low-frequency hearing loss in Japanese. J. Hum. Genet. 52, 510–515 (2007).

44. Bharadwaj, T. et al. The Diverse Genetic Landscape of Hearing Impairment in South African Families. Clin. Genet. 511–520 (2025) doi:10.1111/cge.14765.

45. Porciatti, V. function. 164–170 (2016) doi:10.1016/j.exer.2015.05.008.Electrophysiological.

46. Cameron, A. M., Mahroo, O. A. R. & Lamb, T. D. Dark adaptation of human rod bipolar cells measured from the b -wave of the scotopic electroretinogram. 2, 507–526 (2006).

47. Kakiuchi, D. A. I., Uehara, T., Shiotani, M. & Ito, K. N. Oscillatory potentials in electroretinogram as an early marker of visual abnormalities in vitamin A deficiency. 995–1003 (2015) doi:10.3892/mmr.2014.2852.

48. Ueda, Y., Tammitsu, N., Imai, H., Honda, Y. & Shichida, Y. Recovery of rod-mediated a-wave during light-adaptation in mGluR6-deficient mice. Vision Res. 46, 1655–1664 (2006).

49. Midena, E. et al. Early microvascular and oscillatory potentials changes in human diabetic retina: Amacrine cells and the intraretinal neurovascular crosstalk. J. Clin. Med. 10, (2021).

50. Iv, J. A. B. et al. NIH Public Access. 46, 2540–2551 (2006).

51. Aldunate, E. Z. et al. Conditional Dicer1 depletion using Chrnb4-Cre leads to cone cell death and impaired photopic vision. Sci. Rep. 9, 1–19 (2019).

52. Moskowitz, A., Hansen, R. M., Akula, J. D., Eklund, S. E. & Fultonp, A. B. Rod and rod-driven function in achromatopsia and blue cone monochromatism. Investig. Ophthalmol. Vis. Sci. 50, 950–958 (2009).

53. Angebault, C. et al. ER-mitochondria cross-talk is regulated by the Ca2+ sensor NCS1 and is impaired in Wolfram syndrome. Sci. Signal. 11, (2018).

54. Crouzier, L., et al. Sigma-1 receptor is critical for mitochondrial activity and unfolded protein response in larval zebrafish. Int. J. Mol. Sci. 22, (2021).

55. Fonnum, F., Johnsen, A. & Hassel, B. Use of fluorocitrate and fluoroacetate in the study of brain metabolism. Glia 21, 106–113 (1997).

56. Jassim, A. H. et al. Higher Reliance on Glycolysis Limits Glycolytic Responsiveness in Degenerating Glaucomatous Optic Nerve. Mol. Neurobiol. 56, 7097–7112 (2019).

57. Barboni, P. et al. The Pattern of Retinal Ganglion Cell Loss in Wolfram Syndrome is Distinct From Mitochondrial Optic Neuropathies. Am. J. Ophthalmol. 241, 206–216 (2022).

58. Brandstätter, J. H., Löhrke, S., Morgans, C. W. & Wässle, H. Distributions of two homologous synaptic vesicle proteins, synaptoporin and synaptophysin, in the mammalian retina. J. Comp. Neurol. 370, 1–10 (1996).

59. Balzamino, B. O. et al. Retinal Inflammation and Reactive Müller Cells: Neurotrophins’ Release and Neuroprotective Strategies. Biology (Basel). 13, 1–19 (2024).

60. Cagalinec, M. et al. Role of Mitochondrial Dynamics in Neuronal Development: Mechanism for Wolfram Syndrome. PLoS Biol. 14, 1–28 (2016).

61. Palavicini, J. P. et al. Early disruption of nerve mitochondrial and myelin lipid homeostasis in obesity-induced diabetes. JCI Insight 5, (2020).

62. Zambonin, J. L. et al. Increased mitochondrial content in remyelinated axons : implications for multiple sclerosis. (2011) doi:10.1093/brain/awr110.

63. Bristow, E. A., Griffiths, P. G., Andrews, R. M., Johnson, M. A. & Turnbull, D. M. The distribution of mitochondrial activity in relation to optic nerve structure. Arch. Ophthalmol. 120, 791–796 (2002).

64. Barnes-Vélez, J. A., Aksoy Yasar, F. B. & Hu, J. Myelin lipid metabolism and its role in myelination and myelin maintenance. Innovation 4, 100360 (2023).

65. Tan, S. et al. Pomegranate activates TFEB to promote autophagy-lysosomal fitness and mitophagy. Sci. Rep. 9, 1–18 (2019).

66. Zhuang, X. X. et al. Pharmacological enhancement of TFEB-mediated autophagy alleviated neuronal death in oxidative stress-induced Parkinson’s disease models. Cell Death Dis. 11, (2020).

67. Li, C. H. et al. Spatiotemporal proteomics reveals the biosynthetic lysosomal membrane protein interactome in neurons. Nat. Commun. 15, (2024).

68. Tulva, K. et al. Early trigeminal and sensory impairment and lysosomal dysfunction in accurate models of Wolfram syndrome. Exp. Neurol. 385, (2025).

69. Bora, G. et al. Microtubule-associated protein 1B dysregulates microtubule dynamics and neuronal mitochondrial transport in spinal muscular atrophy. Hum. Mol. Genet. 29, 3935–3944 (2020).

70. Voloboueva, L. A., Suh, S. W., Swanson, R. A. & Giffard, R. G. Inhibition of mitochondrial function in astrocytes: Implications for neuroprotection. J. Neurochem. 102, 1383–1394 (2007).

