## Supplemental data for "WFS1^E864K^ in humans and mice causes Wolfram-like syndrome optic atrophy via early axonal mitochondrial dysfunction"

**Supplemental Material**

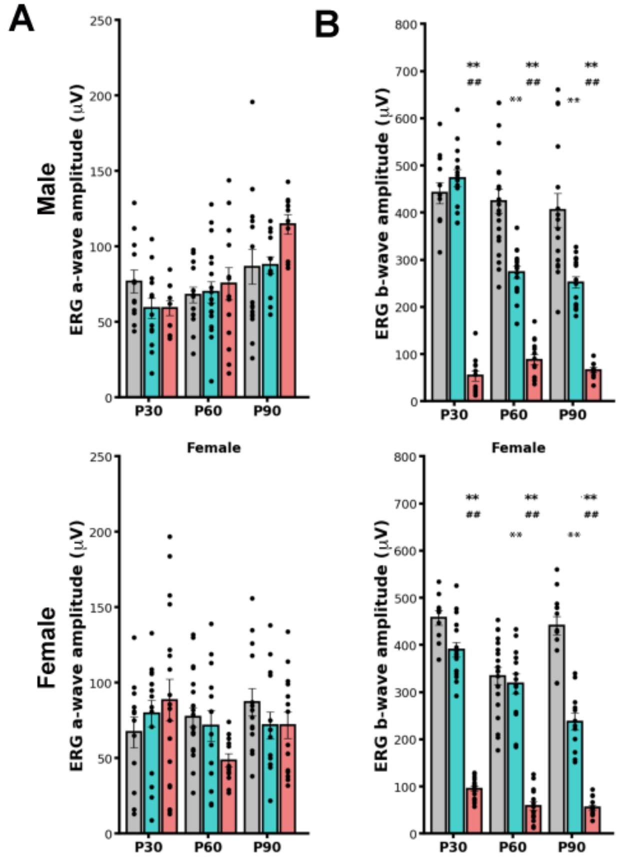


**Supplemental Figure 1. *Wfs1$^{E864K}$* variant induced scotopic retinal disfunction at lower stimulation intensity.**

(A-B) Quantification of a-wave (A) and b-wave (B) amplitude, under scotopic conditions, from *Wfs1$^{WT}$* (grey), *Wfs1$^{E864K/WT}$* (blue) and *Wfs1$^{E864K}$* (red) mice at P30, P60 P90, for both males and females. Data are presented as mean ± SEM from both eyes of 5 to 10 mice. One-way Anova test was performed for each time point, followed by a post-hoc Tukey's test. * $p < 0.05$, ** $p < 0.01$ vs. *Wfs1$^{WT}$* mice, # $p < 0.05$, ## $p < 0.01$ vs. *Wfs1 $^{E864K/ WT}$* mice

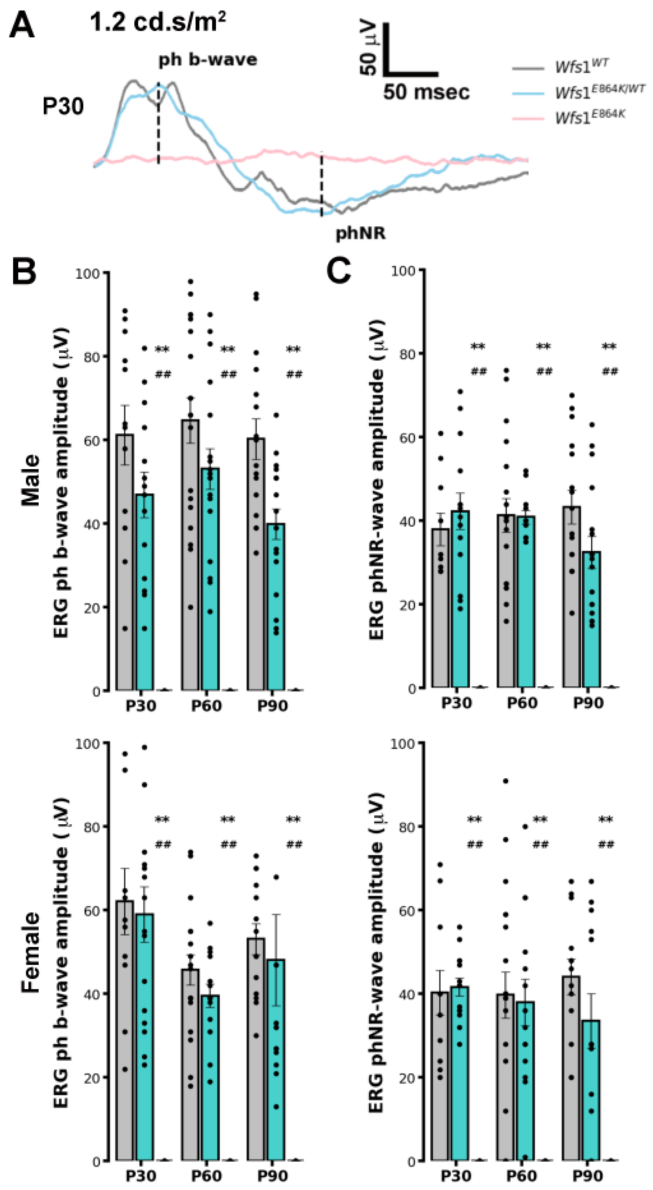


**Supplemental Figure 2. *Wfs1$^{E864K}$* variant induced photopic retinal disfunction.**

(A) Representative photopic traces showing photopic b-wave and photopic negative response (phNR) amplitude from *Wfs1$^{WT}$* (gray), *Wfs1$^{E864K/WT}$* (blue) and *Wfs1$^{E864K}$* (red) male mice at P30. (B - C) Quantification of photopic b-wave (B) and phNR (C) amplitude for *Wfs1$^{WT}$*, *Wfs1$^{E864K/WT}$* and *Wfs1$^{E864K}$* mice at P30, P60 P90, for both males and females. Data are presented as mean ± SEM values from both eyes of 5 to 10 mice. One-way Anova test was performed for each time point, followed by a post-hoc Tukey's test. * $p < 0.05$, ** $p < 0.01$ vs. *Wfs1*$^{WT}$ mice, # $p < 0.05$, ## $p < 0.01$ vs. *Wfs1*$^{E864K/WT}$ mice.

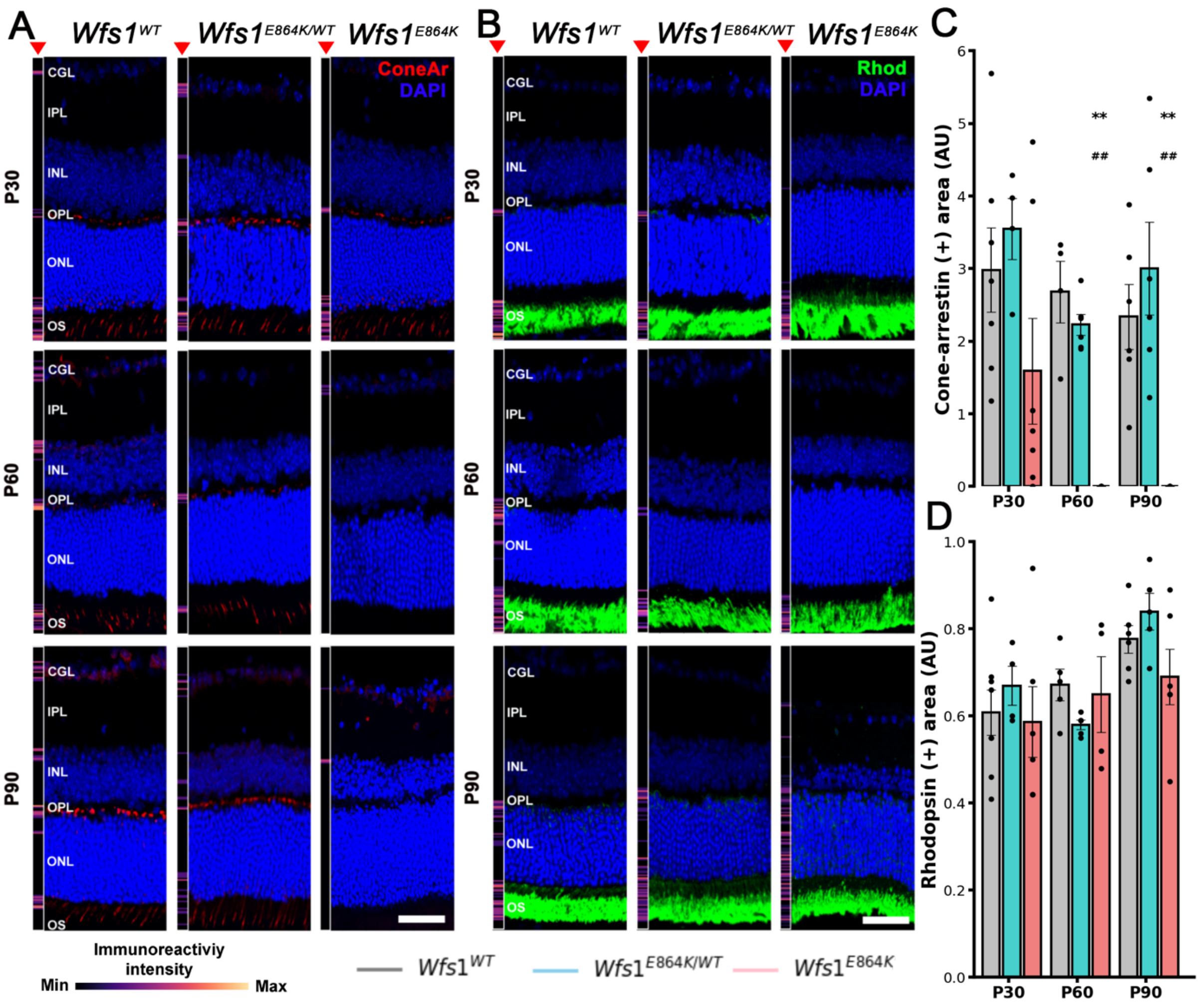


**Supplemental Figure 3. *Wfs1$^{E864K}$* variant induced an alteration in cones but not rods.**

(A - B) Representative images of immunostaining for cone arrestin (A, red) and rhodopsin (Rhod) (B, green) on retina transversal sections, counterstained with DAPI, from *Wfs1$^{WT}$*, *Wfs1$^{E864K/WT}$* and *Wfs1$^{E864K}$* mice at P30, P60 and P90 and heatmap immunoreactive transversal analysis (red head arrow). (C - D) Quantification of cone arrestin (C) and Rhodopsin (D) positive area between *Wfs1$^{WT}$* (gray), *Wfs1$^{E864K/WT}$* (blue) and *Wfs1$^{E864K}$* (red) mice at P30, P60 and P90. Data are presented as mean ± SEM values from 5 to 7 retina sections. Scale bar: 25 mm for all panels in (A) and (B). One-way Anova test was performed for each time point, followed by a post-hoc Tukey's test. ** $p < 0.01$ vs. *Wfs1$^{WT}$* mice, ## $p < 0.01$ vs. *Wfs1$^{E864K/WT}$* mice.

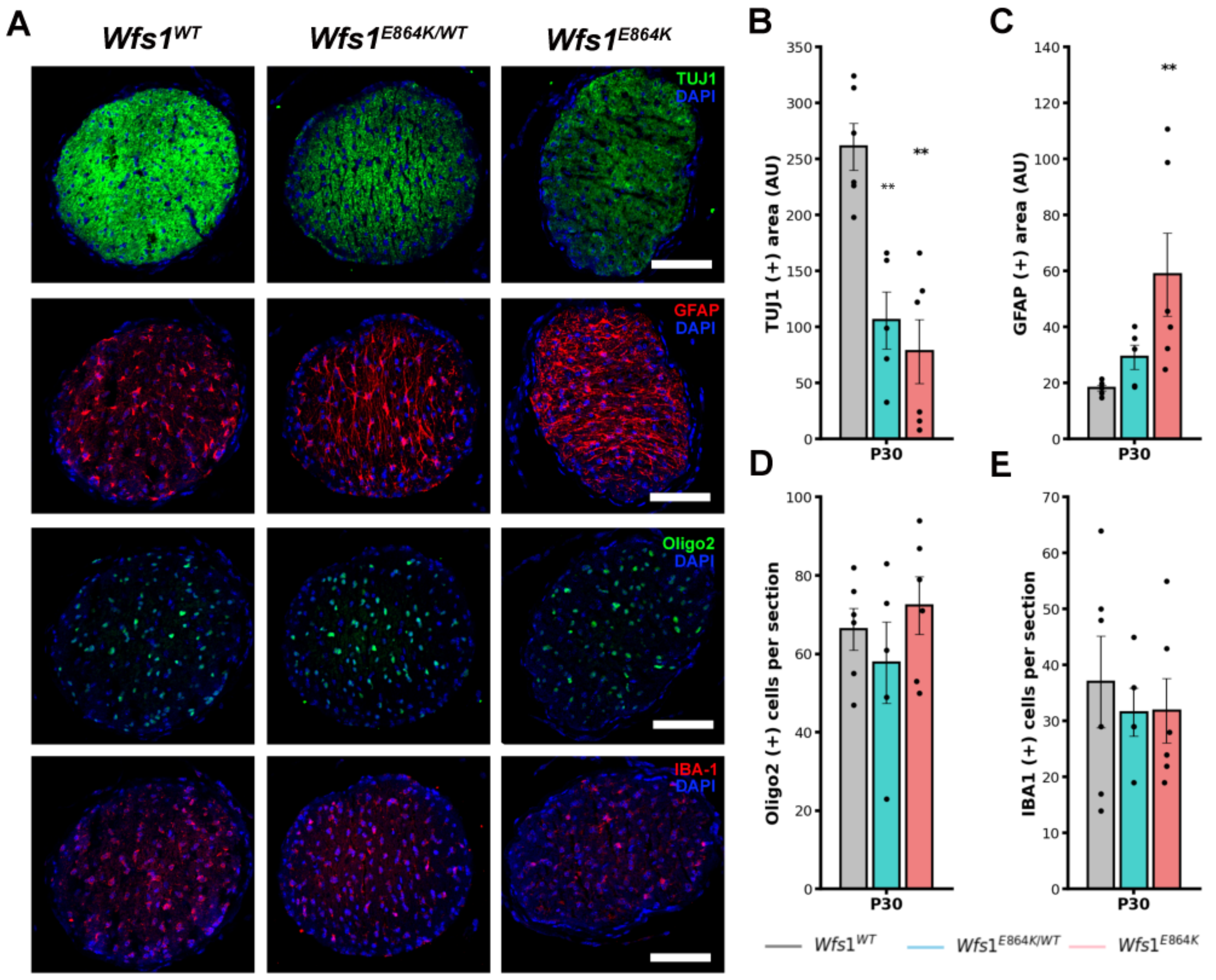


**Supplemental Figure 4.** ***Wfs1$^{E864K}$*** **variant induced an early alteration of axons and reactive gliosis.**

(A) Representative images of immunostaining for TUJ1 (green), GFAP (red), Oligo2 (green) and IBA-1(red) in ON transversal sections, counterstained with DAPI, from *Wfs1$^{WT}$*, *Wfs1$^{E864K/WT}$* and *Wfs1$^{E864K}$* mice at P30. (B - D) Quantification of TUJ1 (B), GFAP (C), IBA-1(D) and Oligo2 (E) positive areas in *Wfs1$^{WT}$* (gray), *Wfs1$^{E864K/WT}$* (blue) and *Wfs1$^{E864K}$* (red) mice. Data are presented as mean ± SEM values from 5 to 7 ONs. Scale bar: 25 mm for all panel in (A). One-way Anova test was performed for each time point, followed by a post-hoc Tukey's test. ** $p < 0.01$ vs. *Wfs1$^{WT}$* mice.

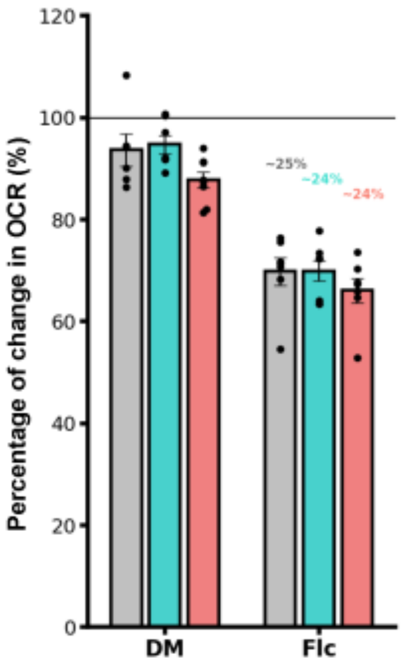


**Supplemental Figure 5. Effect of Fluorocitrate over oxygen consumption rates from ONs.**

Representative graph showing the percentage of change of the basal respiration rate after DMEM (DM) or 2,5 mM of fluorocitrate (Flc) incubation. Data are presented as mean ± SEM values from both ONs of 6 to 8 mice per genotype.

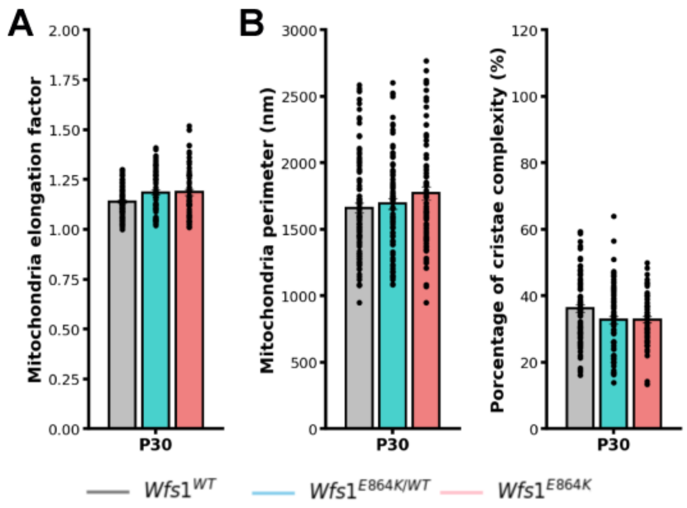


**Supplemental Figure 6. *Wfs1$^{E864K}$* variant did not induce a change in ON mitochondria elongation factor and RGC mitochondria perimeter no cristae complexity.**

(A) Quantification of mitochondria elongation factor in ON mitochondria from *Wfs1$^{WT}$* (gray), *Wfs1$^{E864K/WT}$* (blue) and *Wfs1$^{E864K}$* (red) mice at P30. (B) Quantification of mitochondria perimeter and cristae complexity in RGCS mitochondria from *Wfs1$^{WT}$* (gray), *Wfs1$^{E864K/WT}$* (blue) and *Wfs1$^{E864K}$* (red) mice at P30. Data are presented as mean ± SEM values from 5 to 6 ONs. One-way Anova test was performed for each time point.

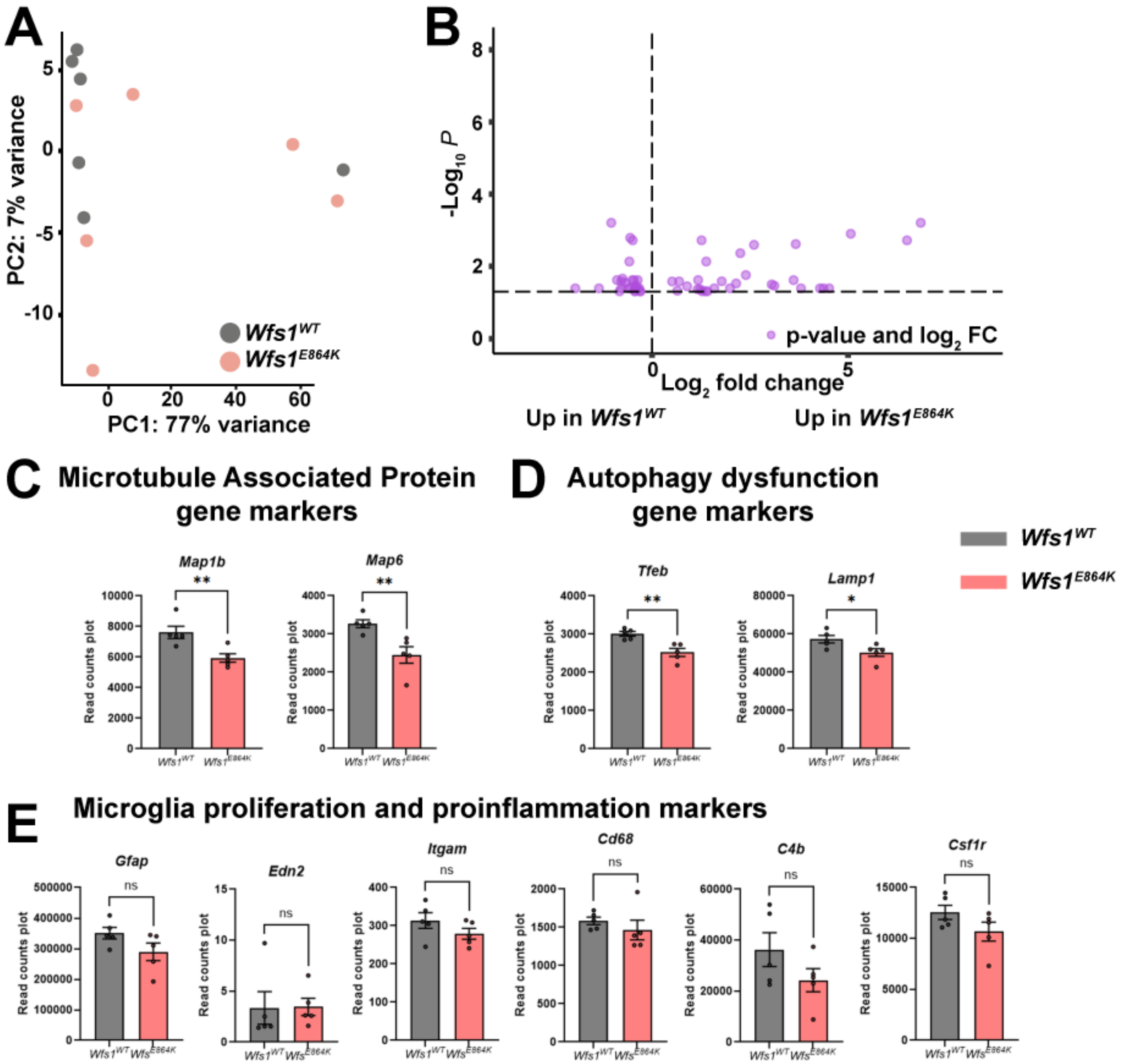


**Supplemental Figure 7. RNA-seq analysis of *Wfs1$^{WT}$* and *Wfs1$^{E864K}$* optic nerves.** (A) Principal component analysis (PCA) plot of *Wfs1$^{WT}$* and *Wfs1$^{E864K}$* RNA-seq analysis illustrating the distribution of gene expression of the samples (n=5/condition) for both conditions. (B) Volcano plot identifying the differentially expressed genes (DEGs) in the *Wfs1$^{E864K}$* versus *Wfs1$^{WT}$* condition. Purple dots represent the 50 genes that were significantly differentially expressed (padj < 0.05) of which 24 were downregulated and 26 upregulated in the *Wfs1$^{E864K}$* condition. (C-E) RNA-seq-based read count plot gene expression analysis of microtubule associated protein genes

(C), autophagy gene (D) and microglia proliferation and proinflammation genes (E) in $Wfs1^{E864K}$ and $Wfs1^{WT}$ optic nerves.

Unpaired t tests or Mann–Whitney tests were performed between the $Wfs1^{E864K}$ and $Wfs1^{WT}$ condition. *$p < 0.05$, **$p < 0.01$.

| | NZ01 | NL01 |
|---|---|---|
| **Age at last evaluatio** | 10 years old | 17 years old |
| **Gender** | Female | Female |
| **Hearing loss**<br>**Age of onset** | Bilateral, severe SNHL<br>9 years old | Bilateral, severe SNHL<br>6 years old (suspected congenital) |
| **Optic atrophy**<br>**Age of onset** | Bilateral<br>9 years old | Bilateral<br>15 years old |
| **Diabetes *mellitus*** | Not detected | Not detected |
| **Other symptoms** | Developmental delay | Pubertal precox<br>Mild developmental delay |
| **Visual evaluation** | | |
| **Visual acuity** | Affected | Affected |
| **Color vision** | Affected | Normal |
| **Cataract** | Not detected | Not detected |
| **Fundus examination** | Bilateral pale optic disc | Bilateral pale optic disc |
| **OCT examination** | OPL lamination<br>Thinning of RFNL | OPL lamination<br>Thinning of RFNL, RGC and INL |
| **MRI examination** | Reduced optic nerve | No performed |

**Supplemental Table 1.** Data examination of case reported patients with autosomal dominant OA.

SNHL: sensorineural hearing loss; OPL: outer plexiform layer; RFNL: retinal fiber nuclear layer; RGC: retinal ganglion cells; INL: inner nuclear layer; OCT: optical coherence tomography; MRI: magnetic resonance imaging.

| Gene ID | log2 Fold Change | padj |
|---|---|---|
| *Clk1* | 0,51 | 0,0262 |
| *Per1* | 0,64 | 0,0473 |
| *Nrip2* | 0,69 | 0,0261 |
| *Rtp4* | 0,89 | 0,0356 |
| *Isg15* | 1,16 | 0,0431 |
| *Zcchc12* | 1,17 | 0,0240 |
| *Spp1* | 1,22 | 0,0431 |
| *Ccl12* | 1,26 | 0,0019 |
| *Prmt8* | 1,28 | 0,0491 |
| *Scube2* | 1,36 | 0,0487 |
| *Rtl3* | 1,38 | 0,0073 |
| *Gm35850* | 1,41 | 0,0491 |
| *Tdgf1* | 1,59 | 0,0407 |
| *Ramp3* | 1,78 | 0,0260 |
| *Tpsab1* | 2,14 | 0,0297 |
| *Acox2* | 2,25 | 0,0043 |
| *Ano9* | 2,39 | 0,0174 |
| *Arhgap36* | 2,60 | 0,0025 |
| *Stab2* | 3,12 | 0,0343 |
| *Ms4a4d* | 3,67 | 0,0024 |
| *Synpo2l* | 4,29 | 0,0407 |
| *Igkv6-25* | 4,37 | 0,0407 |
| *Myoz3* | 4,52 | 0,0407 |
| *Nkx2-1* | 5,07 | 0,0013 |
| *Bpifa2* | 6,51 | 0,0019 |
| *Oxt* | 6,86 | 0,0006 |

**Supplemental Table 2.** Upregulated genes in *Wfs1*$^{E864K}$ versus *Wfs1*$^{WT}$

| Gene ID | log2 Fold Change | padj |
|---|---|---|
| *Shbg* | -1,96 | 0,0407 |
| *Fat2* | -1,36 | 0,0407 |
| *Igfbpl1* | -1,04 | 0,0006 |
| *Baiap2l1* | -0,90 | 0,0240 |
| *Dhh* | -0,83 | 0,0491 |
| *Pik3r5* | -0,79 | 0,0407 |
| *Bcl3* | -0,79 | 0,0262 |
| *Top2a* | -0,69 | 0,0284 |
| *Rrm2* | -0,61 | 0,0407 |
| *Dclk1* | -0,58 | 0,0073 |
| *Slc35f1* | -0,56 | 0,0016 |
| *Dpysl4* | -0,55 | 0,0407 |
| *Frrs1l* | -0,51 | 0,0240 |
| *Efcc1* | -0,50 | 0,0019 |
| *Lingo1* | -0,46 | 0,0243 |
| *Rita1* | -0,45 | 0,0407 |
| *Srd5a1* | -0,44 | 0,0487 |
| *Dpysl3* | -0,44 | 0,0487 |
| *Pcyox1l* | -0,42 | 0,0343 |
| *Fbxo28* | -0,38 | 0,0407 |
| *Nceh1* | -0,37 | 0,0240 |
| *Rabl3* | -0,33 | 0,0407 |
| *Fdft1* | -0,31 | 0,0491 |
| *Taf6l* | -0,30 | 0,0491 |

**Supplemental Table 3.** Downregulated genes in $Wfs1^{E864K}$ versus $Wfs1^{WT}$

| Gene ID | log2 Fold Change | pvalue | padj |
|---|---|---|---|
| *Map1b* | -0,36 | 0,0019 | 0,1812 |
| *Map6* | -0,42 | 0,0054 | 0,2962 |
| *Tfeb* | -0,25 | 0,0042 | 0,2630 |
| *Lamp1* | -0,19 | 0,0503 | 0,5686 |

**Supplemental Table 4.** Microtubules associated protein and autophagy dysfunction gene markers downregulated in *Wfs1*$^{E864K}$ versus *Wfs1*$^{WT}$
